# Uncovering the Genetic Profiles Underlying the Intrinsic Organization of the Human Cerebellum

**DOI:** 10.1101/2021.06.23.448673

**Authors:** Yaping Wang, Lin Chai, Congying Chu, Deying Li, Chaohong Gao, Xia Wu, Zhengyi Yang, Yu Zhang, Junhai Xu, Jens Randel Nyengaard, Simon B. Eickhoff, Bing Liu, Kristoffer Hougaard Madsen, Tianzi Jiang, Lingzhong Fan

## Abstract

The functional diversity of the human cerebellum is largely believed to be derived more from its extensive connections rather than being limited to its mostly invariant architecture. However, whether and how the determination of cerebellar connections in its intrinsic organization interact with microscale gene expression is still unknown. Here we decode the genetic profiles of the cerebellar functional organization by investigating the genetic substrates simultaneously linking cerebellar functional heterogeneity and its drivers, i.e., the connections. We not only identified 443 network-specific genes but also discovered that their co-expression pattern correlated strongly with intra-cerebellar functional connectivity (FC). Ninety of these genes were also linked to the FC of cortico-cerebellar cognitive-limbic networks. To further discover the biological functions of these genes, we performed a “virtual gene knock-out” by observing the change in the coupling between gene co-expression and FC and divided the genes into two subsets, i.e., a positive gene contribution indicator (GCI^+^) involved in cerebellar neurodevelopment and a negative gene set (GCI^−^) related to neurotransmission. A more interesting finding is that GCI^−^ is significantly linked with the cerebellar connectivity-behavior association and many recognized brain diseases that are closely linked with the cerebellar functional abnormalities. Our results could collectively help to rethink the genetic substrates underlying the cerebellar functional organization and offer possible micro-macro interacted mechanistic interpretations of the cerebellum-involved high order functions and dysfunctions in neuropsychiatric disorders.

## Introduction

Converging evidence from animal and human studies is advancing our understanding of the human cerebellum, which is engaged in motor, complex cognitive, and emotional behaviors [1–4]. While such functional diversity of the cerebellum was believed to derive more from its extensive afferent and efferent connections to extra-cerebellar structures, rather than being limited to a regular lattice-like anatomic feature of cerebellar-cortical cytoarchitecture [1, 5–8]. Considering the widely accepted understanding that the macroscale functional organization of the human nervous system is ultimately regulated by the underlying microscale gene expression [9–12], it is thus intriguing to unravel the genetic profiles underlying the cerebellar functional organization. As yet, what remains unclear is whether and how the hypothesized determination of cerebellar connections in its intrinsic functional organization interact with microscale gene expression.

To date, the genetic mechanism supporting the functional organization of the human cerebellum is mainly unknown. Only a few studies have attempted to investigate the gene expression pattern of the human cerebellum, but they provided inconsistent results in gene expression variability. For instance, Hawrylycz et al. [13] and Negi and Guda [14] both found that gene expression is highly homogeneous across the anatomical regions of the healthy adult cerebellum. In contrast, Aldinger et al. [15] and Wang and Zoghbi [12] found that cerebellar development and function are governed by the precise regulation of molecular and cellular programs and that the gene expression pattern is heterogeneous across spatial and temporal scales. In addition, differences in gene expression patterns between the cerebellar gyri and sulci [16], and considerable cerebellar regional specializations containing specific cell types, as revealed by high-throughput single-nucleus RNA-seq [17] have been found in the mouse cerebellum. Because relevant studies that showed homogeneity [13, 14] explored the overall cerebellar genetic expression pattern across its gross macro-anatomical boundaries (e.g., cerebellar lobules) and might have failed to fully reflect the functional organization of the human cerebellum [18, 19]. This inconsistency in the genetic variability of the cerebellum needs to be further explored. In the past decade, functional topological maps describing the organization of the human cerebellum using task [20] and task-free functional magnetic resonance imaging (fMRI) [21, 22], specifically, separate cerebellar functional networks [21, 22] and intra-cerebellar functional gradients [23], have been proposed. In particular, Buckner et al. [21] employed resting-state functional connectivity (FC) of the cerebello-cortical circuit as a tool to map the intrinsic functional architecture of the human cerebellum and proposed a possible functional parcellation into 7 networks and 17 networks. It is thus possible to decode the genetic profiles of the cerebellar functional organization by investigating the molecular genetic substrates linking cerebellar functional heterogeneity and its drivers, i.e., the connections. One promising approach is imaging-transcriptomics analysis [24–26], which allows the brain-wide spatial analysis of microscopic transcriptome data combined with macroscopic neuroimaging phenotypes [9]. Moreover, it also offers the opportunity to link the transcriptome data with the behavior variations via the neuroimaging [27]. These cross-scale analyses could provide a better understanding of the interaction between microscale gene expression and macroscale functional network, and ultimately involved in the individual behaviors, and importantly, their putative multiscale interactions in cerebellar related diseases [28].

Thus, our goal was to investigate the neurobiological genetic substrates underlying the functional organization of the human cerebellum. More specifically, we will seek to address the following three progressive questions (Fig. S1). Are there differentially expressed genes in the diverse intra-cerebellar functional networks with largely invariant cytoarchitecture? How do gene expression and cerebellar connectivity relate to each other, and is there any link between this association and human behavior, as well as brain disease? In the current study, we first examined network-specific genes by differential gene expression analysis across diverse cerebellar functional networks characterized by largely invariant cytoarchitecture. This analysis allowed us to further investigate the genetic explanations to the cerebellar inconsistency between functional heterogeneity and cytoarchitecture near-homogeneity. Then to explore the relationship between these network-specific genes with the cerebellar connection, we constructed the gene co-expression matrix using the network-specific genes and found it is highly correlated with both intra-cerebellar and cerebello-cortical cognitive-limbic FC. Furthermore, we tested the contribution of each network-specific gene to this correlation by virtual gene knock-out (KO) and divided the network-specific genes into two subsets, i.e., a positive gene contribution indicator (GCI^+^) and a negative gene set (GCI^−^). We found that the GCI^+^ appears to be mainly involved in cerebellar neurodevelopment. Whereas GCI^−^ seems to be related to neurotransmission, emotion-cognitional behaviors and is significantly enriched in various neuropsychiatric disorders that are closely linked with cerebellar functional abnormalities. Together, the current exploration provides a starting point for associating the genetic and behavior markers of the functional network to cerebellar involvements in higher order non-motor functions and dysfunctions in various neuropsychiatric disorders [1, 29, 30].

## Materials and Methods

The schematic of the experimental design is depicted in Fig. 1 and includes three steps. Step 1 (Fig. 1A): to investigate whether network-specific genes occurred across cerebellar functional networks, we performed the differential gene expression analysis based on the combination of the Allen Human Brain Atlas (AHBA) transcriptome data [9] with a cerebellar functional parcellation atlas [21]. Step 2 (Fig. 1B): then, the co-expression matrix was constructed using the network-specific genes from step 1 and compared with FC to explore their overall correlation (referred to as the Gene-FC correlation for simplicity). Meanwhile, the relationship between gene and cerebello-cortical FC also has been explored. Step 3 (Fig. 1C): the “virtual gene knock-out” was leveraged to examine the direction of each gene’s contribution to this Gene-FC correlation from step 2 and used to separate the network-specific genes into two subsets. Furthermore, we applied a series of functional annotation tools to explore the role of these genes, including gene enrichment analysis, Behavior-FC-Gene mapping analysis, disease enrichment analysis, and integrative temporal specificity analysis.

**Fig. 1.**
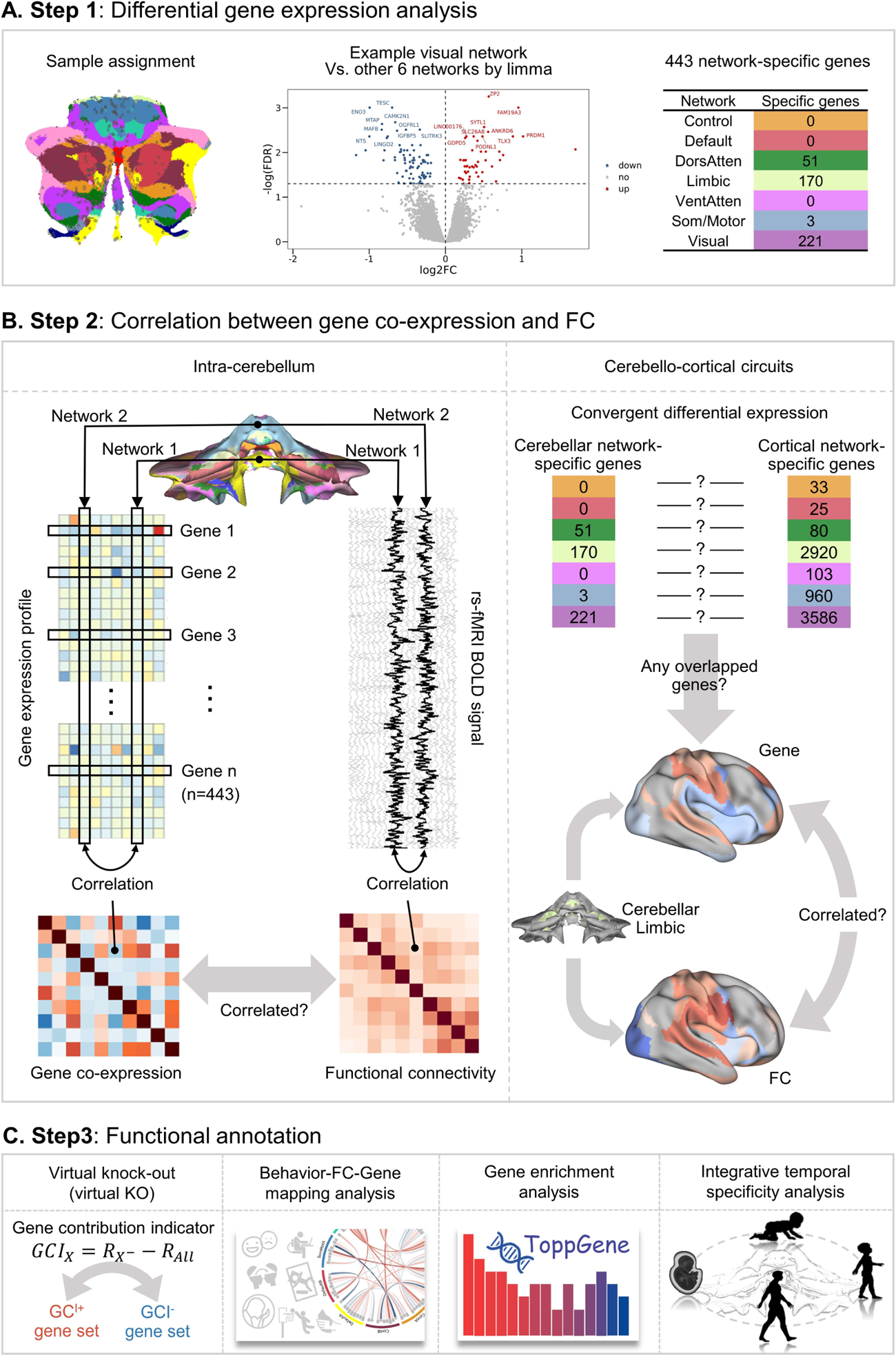
Analysis pipeline. **A Step 1**: differential gene expression analysis. We assigned the AHBA cerebellar samples into seven cerebellar functional networks (left) [21] and averaged each gene’s expression within the same network individually. Then we compared the gene expression in each network with all the other networks by limma [31] (middle) with a fold change > 1 and *p* < 0.05 (FDR corrected) as an indicator (Red indicates that the genes were found significantly positively expressed in the visual network.). Thus, we obtained the network-specific genes for the seven networks (right). **B Step 2:** correlations between the gene co-expression and the FC included intra-cerebellar and cerebello-cortical circuit. Intra-cerebellum: for each pair of networks, we calculated the gene expression similarity between them using the cerebellar network-specific genes and then constructed the gene co-expression matrix. The FC matrix was constructed by correlating the BOLD signal for all pairs. Then the relationship between the genetic correlation and functional correlation was evaluated. Cerebello-cortical circuit: we first defined the cortical network-specific genes as we did for the cerebellum and tested whether any convergently differentially expressed genes occurred. Then we used the overlapping genes to obtain the cortical genetic correlation for each cerebellar network and evaluated the relationship between the cortical genetic and functional correlation for each cerebellar network. **C Step 3:** functional annotation includes virtual gene knock-out (KO), Behavior-FC-Gene mapping, gene enrichment analysis, and integrative temporal specificity analysis.

### AHBA preprocessing

The AHBA [9] is a publicly available transcriptome dataset (http://www.brain-map.org), which provides normalized microarray gene expression data from six adult donors (ages 24, 31, 34, 49, 55, and 57 years old; *n* = 4 left-hemisphere only, *n* = 2 both left and right hemispheres). Table S1 shows the demographic information. Tissue collection was approved by Institutional Review Boards of the Maryland Department of Health and Hygiene, University of Maryland Baltimore and University of California Irvine, and the informed consents were obtained from decedent next-of-kin [9]. Although AHBA provides gene expression from only six adult donors, it still has many unprecedented advantages. While some existing human gene expression atlases just cover multiple brain regions, only the AHBA delivers high-resolution coverage of nearly the entire brain [32]. It includes the expression of more than 20,000 genes taken from 3,702 spatially distinct tissue samples [9] ranging from the cerebral cortex to the cerebellum.

Referred to Anderson et al. [26], the current preprocessing pipeline included data filtering, probe selection, sample selection, and assignment. We first filtered the probes with the AHBA binary indicator to mitigate the background noise and excluded probes without an Entrez ID. Then for the genes that corresponded to two or more probes, we chose the probe with the maximum summed adjacency to represent the corresponding gene expression; otherwise, we retained the probe with the highest mean expression, using the CollapseRows function [33] in R. The first two steps generated 20,738 unique mRNA probes, which provided expression data for 20,738 genes. As suggested by Arnatkeviciute et al. [32] and given the known transcriptional differences [13] between the cortical and sub-cortical regions and the cerebellum, we separated the cortical and cerebellar samples a priori based on the slab type and structure name provided by AHBA and processed them separately later. In the end, 337 samples were retained for the cerebellar cortex and 1,701 samples for the cortical cortex.

Finally, we respectively assigned these 337 cerebellum samples and 1,701 cortical samples into the cerebellar functional networks atlas [21] and cortical functional networks atlas [34], both of which have 7- and 17-network parcellation strategies. For each cerebellar sample, we first generated a single 1 × 1 × 1 mm^3^ region of interest (ROI) at the MNI coordinate for each sample using the AFNI 3dmaskdump -nbox function [35]. The network label from either 7- or 17-network parcellation was assigned if the ROI fell within a cerebellar network of the Buckner atlas. Considering the uneven and discrete sampling of the AHBA data [9], if the 1 × 1 × 1 mm^3^ ROI did not overlap with any network, the associated ROI was expanded to 3 × 3 × 3 mm^3^. And if the 3 × 3 × 3 mm^3^ ROI overlapped with the functional atlas, the network that had the maximum number of shared voxels with the ROI was assigned. Otherwise, the steps above were repeated for a 5 × 5 × 5 mm^3^ ROI. The cerebellar samples were excluded if the 5 × 5 × 5 mm^3^ ROI did not overlap with any cerebellar networks. Tables S2 and S3 show the distributions of the cerebellar sample assignment for the 7-network and 17-network atlases. The assignment of the AHBA cortical samples into the cortical functional network atlas was consistent with the method used for the cerebellum, and the cortical sample distributions are shown in Tables S4 and S5.

### Differential gene expression analysis across functional networks

The gene expressions of the cerebellar samples within the same network were averaged for each gene across the samples individually, resulting in 20,738 genes × 7 or 17 networks matrices for each donor. Here we used as many of the most original overall genes as possible, because some existing methods [13, 32] for screening genes do not consider the specificity of gene expression within the human cerebellum. Then we calculated the differentially expressed genes across the 7 networks using the R limma package [31] by comparing the gene expression in one network (e.g., control) with the remaining 6 networks (e.g., default, limbic, visual, etc.). Since the gene expression vectors of the same network derived from different donors can be treated as biological replicates [26], the limma’s duplicatedCorrelation tool [36] was leveraged to evaluate the independence between replicates. Specifically, the correlations between replicates were estimated using the duplicateCorrelation function, which individually fits a mixed linear model by restricted maximum likelihood for each gene and returns a consensus correlation [36]. This consensus correlation between replicates would then be incorporated into the limma’s linear model to preserve more information than simply averaging the biological replicates [36]. The traditional minimum fold change threshold was unsuitable for determining biologically meaningful but subtly different expressions [37], in particular for the cerebellum, due to the highly homogenous cytoarchitecture. Instead, we applied the Benjamini-Hochberg (BH) method to control the false discovery rate (FDR), and the statistical threshold *p* ≤ 0.05 (FDR corrected) combined with a fold change > 1 was used as the key indicator for differentially expressed genes. For simplicity, the differentially expressed genes across cerebellar networks are referred to as cerebellar network-specific genes throughout this paper. The cortical network-specific genes were identified in the same way. The only difference was that the gene expression of the cortical samples was first averaged within each parcel (51 and 114 parcels, which corresponded to the 7- and 17-networks, respectively) [34] and then averaged within each network.

To test the specificity of these network-specific genes derived based on task-free 7 functional networks, we also applied the differential gene expression analysis with the same procedure in other different cerebellar atlases. There are the different resolution of the same task-free functional atlas [21], i.e., 17-network parcellation, independent task-based multi-domain task battery (MDTB) [20] functional atlas and different resolutions, i.e., 11-, 28-lobular parcellations, of the anatomical atlas [38]. Then we estimated the overlapping genes between the network-specific genes derived based on the task-free 7 functional networks with other atlases. Moreover, we applied the differential gene expression analysis in only 4 left-hemisphere donors with the same procedure to test the sensibility of these network-specific genes.

### Cerebellar resting-state functional connectivity (FC)

The minimally preprocessed [39, 40] Human Connectome Project (HCP) S1200 release dataset [41], which has 1,018 subjects (aged from 22 to 37 years old) with both structural MRI and resting-state functional MRI (rs-fMRI, HCP S1200 manual), was used. The preprocessing pipeline includes artifact correction (correction of gradient nonlinearity distortion, realignment for head motion, registration of fMRI data using structural data, reduction of geometric distortions due to B0 field inhomogeneity, etc.) as well as denoising by ICA-FIX [42, 43]. Time courses were extracted from these CIFTI grayordinate-format preprocessed rs-fMRI images, and the global signal was regressed as well. The resting-state blood oxygen level-dependent time series were averaged within each cortical parcel of the 7- or 17-network cortical atlases and within each cerebellar network of the 7- or 17-network cerebellar atlases [21], separately. The FC within the cerebellum was computed using Pearson’s correlation for the averaged time courses for each interest ROI. Because four runs were performed for each subject, the correlation values were separately calculated for each run, Fisher’s z-transformed, and averaged across the runs, resulting in a 17 × 17 cerebellar networks matrix. The same process was used to calculate the correlations between each functional cerebellar network and each cortical parcel, resulting in a 114 cortical parcels × 7 cerebellar networks functional correlation matrix, which represents the FC across the cerebello-cortical circuit. Regardless of whether the FC was within the cerebellum or across the cerebello-cortical circuit, both categories of FC were defined using the more fine-grained 17-network parcellation to increase the spatial resolution. The only exception was that the cerebellar 7-network was applied while calculating the FC across the cerebello-cortical circuit to compare each cerebellar network directly.

### Correlation between gene co-expression with intra-cerebellar FC

To fully capture the genetic correlation with the FC within the cerebellum, we leveraged the genetic samples of the two bi-hemisphere donors when constructing the gene co-expression matrix since the FC of the cerebellum is bilateral. Therefore, the gene co-expression was analyzed for the two bi-hemisphere donors using the network-specific genes derived from all six donors across 7 networks, using a finer 17-network parcellation to increase the spatial resolution. Ten networks that contained samples from both bi-hemisphere donors were retained (Table S3). For each bi-hemispheric donor, the log2 gene expression of the cerebellar samples was mean-normalized and then averaged within each network. The cerebellar 10 × 10 networks correlation matrix was calculated using Spearman’s correlations individually (since the non-normality of gene expression data), then Fisher’s z-transformed, and finally averaged to construct the final 10 networks gene co-expression matrix. The correlation significance level of the gene co-expression was evaluated using the overlap between the correlation significance matrix for these two individuals after being adjusted by Bonferroni correction. Meanwhile, we transformed the 17 × 17 networks FC matrix into a 10 × 10 networks size to be consistent with the gene co-expression matrix. Finally, the relationship between the 10 × 10 networks gene co-expression and the 10 × 10 networks FC matrix was computed using Pearson’s correlation. The correlation between the gene co-expression and FC is referred to as the Gene-FC correlation throughout the present paper for simplicity.

To test the significance of the Gene-FC relationships while taking into account the spatial autocorrelation (SA), we leveraged Brain Surrogate Maps with Autocorrelated Spatial Heterogeneity (BrainSMASH) [44] to generate 10,000 surrogate maps which persevered the genetic SA of the 443 genes × 106 samples matrix. The 106 samples were equivalent to the amount of samples for the 10 networks that were used to construct the gene co-expression matrix since they contained samples from both bi-hemisphere donors. The parameter “knn” was set to 100 to keep it roughly consistent with the number of samples. These 10,000 surrogate maps were used to construct the gene co-expression matrix and thus generate an empirical null distribution of the Gene-FC correlation while taking SA into account. The *p* value (p_SA_) was defined as the proportion of correlation values produced by the surrogate maps that exceeded the correlation coefficient for the real data.

In addition, to evaluate the robustness of the verified Gene-FC correlation within the cerebellum, we also recalculated it using several different parcellations, i.e., task-free 7-network parcellation, and independent task-based MDTB functional parcellation [20]. The criteria for each step were consistent with our primary method. Moreover, we employed a control test to learn whether the Gene-FC correlation could be obtained using only the network-specific genes. That is, no Gene-FC correlation while using other genes. We randomly selected 443 genes from the full gene set without the network-specific genes and referred to them as non-network-specific genes. Then we calculated the Gene-FC correlation using the non-network-specific genes and ran this step randomly 10,000 times. In addition to these thresholdless non-network-specific genes, we applied a set of thresholds to the averaged original log2 gene expression data to confirm that these non-network-specific genes were expressed in the cerebellum. Doing this can also help us test whether the gene co-expression pattern constructed using these threshold non-network-specific genes was correlated with FC.

Lastly, we also verified the robustness of this Gene-FC correlation using the 218 unrelated participants from the HCP S1200 release [41]. Besides, to constitute an independent neuroimaging validation dataset, we leveraged 296 participants with four runs (aged 38−58 years old) from preprocessed HCP-Aging Lifespan 2.0 Release [45]. This dataset, together with the 1,018 participants from the HCP S1200 release of our primary strategy, neatly covered the age ranges of AHBA (extends from 24 to 57 years old). The analysis strategy for these two additional datasets is similar to our primary approach.

### Correlation between gene co-expression and FC across the cerebello-cortical circuit

To thoroughly investigate the cerebellar functional organization, we also explored the relationship between the cerebello-cortical FC and the genetic correlation based on the strategy used in Anderson et al. [26]. First, we defined the network-specific genes in the cortex using the same procedure as we had for the cerebellum and examined the genes that overlapped within the same network of the cerebellum and the cortex. Then the gene co-expression matrix was constructed between 6 cerebellar networks and 59 cortical parcels from the two bi-hemisphere donors, using the 90 unique genes derived from the overlap between the cortical network-specific genes and the cerebellar network-specific genes. Here, the cerebellar 7-network parcellation was selected to compare the different cerebellar networks directly. The visual network was excluded because it only had two samples from one of the 2 bi-hemisphere donors. For the cerebral cortex, 59 cortical parcels that contained samples from both bi-hemisphere donors were estimated. The log2 mean-normalized expression within each cerebellar network and each cortical parcel was estimated individually and correlated using Spearman’s *ρ*, Fisher’s z-transformed, and averaged. We transformed the 114 cortical parcels × 7 cerebellar networks FC matrix into 59 cortical parcels × 6 cerebellar networks size to be consistent with the gene co-expression matrix. Finally, the relationship between the cortical genetic correlation and the cerebello-cortical FC matrix was computed using Pearson’s correlation across six cerebellar networks and adjusted by the BH method to correct for multiple comparisons.

### Gene functional annotation

#### Virtual gene knock-out (KO)

To extend our investigation of the overall relationship between gene co-expression and FC within the cerebellum, we referred to a similar previous approach [46, 47] and termed it the “Virtual Gene Knock-out (KO)” to evaluate each gene’s contribution to the Gene-FC correlation. In brief, we deleted each of the 443 cerebellar network-specific genes one-by-one to simulate the gene KO, then constructed the gene co-expression matrix without that gene, analyzed the correlation between the FC and the gene co-expression, and finally calculated the difference in the correlation coefficient between before and after the simulated deletion, with the result being defined as the gene contribution indicator (GCI) [46]. Based on the GCI, we identified two different gene sets that had opposite effects on the Gene-FC correlation: a GCI positive gene set (GCI^+^) and a GCI negative gene set (GCI^−^). The virtual KO of GCI^+^ increased the Gene-FC correlation, and, accordingly, its expression decreased the Gene-FC correlation; in contrast, the virtual KO of GCI^−^ decreased the correlation, and, accordingly, its expression increased the Gene-FC correlation.

#### Behavior-FC-Gene mapping analysis

To evaluate the relationship between these genes and human behaviors, we leveraged the stable brain-behavior space strategy proposed by Ji et al. [27], which could be briefly divided into the following three sections. These procedures were called Behavioral-FC-Gene mapping analysis to facilitate a better understanding and fluency of the paper.

1. Principal component analysis (PCA) of HCP behavioral data: 59 behavioral measures [48] (Table S6) covering six behavior categories that represent the general domains of human behavior were used, including alertness, cognition, emotion, motor, personality, and sensory. The 218 HCP unrelated participants were selected to avoid violation of the exchangeability assumption of permutation tests, and subjects with missing data were excluded, resulting in 211 participants. The significant principal components (PCs) were calculated by a permutation test, which randomly and independently shuffled subjects’ order for each behavior measure to re-run PCA 10,000 times to establish the null model. The PCs that explained variance exceeded chance (*p* < 0.05 across permutation test) were considered significant and retained for the next step. The nomination of PCs was based on the pattern of loadings on the original 59 behavior measures.
2. Mass univariate Behavior-FC map: the relationship between significant PCs scores and individual intra-cerebellar FC between 17 networks was quantified by a mass univariate regression procedure called PALM tool [49] across 211 unrelated participants. For each significant PC, the derived regression coefficients on the total 136 functional connections were then Z-scored and referred to as the Behavior-FC map for this PC. The significances of the regression coefficients were assessed by a permutation test, which randomly shuffled the order of the subjects for each significant PC score 10,000 times.
3. Behavior-FC-Gene co-expression mapping analysis: to evaluate the relationship between the Behavior-FC map and the co-expression map of GCI^+^ and GCI^−^, we first constructed these maps across the ten cerebellar networks that corresponded to the networks containing genetic samples from both AHBA bi-hemisphere donors. The relationship between the Behavior-FC maps of eight significant PCs and the co-expression patterns of GCI^+^ and GCI^−^ was calculated by the Pearson’s correlation respectively, and the significance was assessed by a permutation test which randomly shuffled the gene co-expression pattern 10,000 times.

#### GO, pathway, and disorder enrichment analysis (ToppGene portal)

To characterize the biological role of GCI^+^ and GCI^−^, we applied the *ToppGene portal* [50] to conduct a gene ontology (GO), pathway, and disorder enrichment analysis. The GO [51] enrichment analysis provides ontologies to describe accumulated knowledge of genes in three biological domains: biological process, cellular component, and molecular function. The BH method for FDR (FDR-BH correction) (*p* < 0.05) was chosen to correct for multiple comparisons.

#### Integrative temporal specificity analysis

To investigate the overall temporal expression features of these genes, we applied an online cell type-specific expression analysis (CSEA) tool [52] to do the enrichment analysis of the genes within the cerebellum during different lifespan windows. Here, a specificity index probability (pSI = 0.05, 0.01, 0.001, and 0.0001, permutation corrected) was used to define the probability of a gene being expressed in each time window relative to all other time windows to represent the varying stringencies for enrichment. The significance of the overlap between the interest gene set and those enriched in a specific time window was evaluated by Fisher’s exact test, and the BH method for FDR (FDR-BH correction) was chosen to correct for multiple comparisons.

### Data and availability

R 3.6.1 and custom scripts were used to perform statistical analysis. All R packages were mentioned explicitly in the text where the package was used. The code is freely available at https://github.com/FANLabCASIA/CerebellarGeneFCCorrelation. The ToppGene (https://toppgene.cchmc.org) and CSEA tool (http://genetics.wustl.edu/jdlab/csea-tool-2/) which used to do the functional annotation of genes were all freely accessible. All data needed to evaluate the conclusions in this study are present in the article and the Supplementary Materials.

## Results

### The cerebellar network-specific genes derived based on the functional segregation within the cerebellum

The genes that were expressed much more in one network than in all the other six networks in the cerebellum and cerebral cortex were identified based on the differential gene expression analysis and are referred to as cerebellar network-specific genes and cortical network-specific genes, respectively. We identified 443 cerebellar network-specific genes (Supplementary Sheets 1 and 3) using all samples from six donors across 7 networks. The distribution of these network-specific genes is shown in Table 1 (Supplementary Sheet 2), which shows that these were mainly expressed in the limbic (*n* = 170), dorsal attention (*n* = 51), somatomotor (*n* = 3), and visual (*n* = 221) networks.

**Table 1.** Counts of significant differentially expressed genes within each network compared to other networks, referred to as the network-specific genes. The cerebellar (*n* = 443, left column, Supplementary Sheets 2 and 3) and cortical network-specific genes (*n* = 6987, middle column, Supplementary Sheets 7 and 8) were defined across the cerebellar [21] and cortical [34] 7-network strategies, respectively. The rightmost column measures the overlap between the cerebellar and cortical network-specific genes for each network (Supplementary Sheets 9 and 10).

| Networks | Cerebellum | Cortex | Overlapping genes |
| --- | --- | --- | --- |
| Control | 0 | 33 | 0 |
| Default | 0 | 25 | 0 |
| Dorsal Attention | 51 | 80 | 0 |
| Limbic | 170 | 2920 | 56 |
| Ventral Attention | 0 | 103 | 0 |
| SomatoMotor | 3 | 960 | 2 |
| Visual | 221 | 3586 | 33 |
| Total (unique) | 443 | 6987 | 90 |

To test the specificity of these 443 network-specific genes in reflecting the difference of gene expression across task-free cerebellar functional atlas, we applied the differential expression analysis with the same procedure in other different cerebellar atlases (see more details in “Methods”). More overlaps between these 443 network-specific genes with the differentially expressed genes were observed in the different resolution of the same task-free functional atlas, whereas less in the independent task-based functional atlas and different resolutions of the lobular atlas (Supplementary Sheet 4). We also tested the sensitivity of network-specific genes in only four left-hemisphere donors with the same procedure and observed 230 cerebellar network-specific genes (Supplementary Sheet 5) with preferentially expressed in the limbic (*n* = 49, overlap = 44) and visual (*n* = 181, overlap = 97) networks.

Meanwhile, we obtained 6,987 cortical network-specific genes (Supplementary Sheets 6−8 and Table 1) using the same strategy and found that the cerebellar and cortical network-specific genes distribution patterns across 7-network parcellation were highly correlated (*r* = 0.95, *p* = 0.001). Moreover, we found that 90 of these 443 cerebellar network-specific genes (∼20%) (Supplementary Sheets 9 and 10 and Table 1) were convergently expressed in the cerebral cortex (overlap in limbic = 56, somatomotor = 26, visual = 33). These results mean that the 56 limbic genes were differentially expressed in the limbic cortex and the limbic cerebellum and that the two somatomotor genes were differentially expressed in the somatomotor cortex and somatomotor cerebellum, as well as the 33 overlap genes in the visual network.

### The co-expression of the cerebellar network-specific genes highly correlated with intra-cerebellar FC

Using the 443 cerebellar network-specific genes, we constructed the gene co-expression matrix for the two bi-hemisphere donors and explored the relationship between gene correlation and FC within the cerebellum. Noted, the 443 network-specific genes were defined based on 7-nework parcellation, while the genetic co-expression was constructed using 17-network parcellation to increase the spatial precision. Across all the available network-network pairs, the genetic co-expression correlates with the FC within the cerebellum (*r* = 0.48, *p*_SA_ = 0.008, Fig. 2), which remains significant when assessed by the spatial autocorrelation (SA) preserving strategy. This correlation between gene co-expression and FC was referred to as Gene-FC correlation throughout the present paper for simplicity. To validate the Gene-FC correlation within the cerebellum, we also leveraged the different resolution of the same task-free functional atlas and the independent task-based MDTB functional parcellation [20] to re-perform the aforementioned steps (Fig. 2D). The gene co-expression and FC within the cerebellum also correlated when analyzed based on the 7-network parcellation (Figs. 2D and S2): *r* = 0.76, *p*_SA_ = 0.006, and the MDTB functional parcellation (Figs. 2D and S3): *r* = 0.42, *p*_SA_ < 0.001. Moreover, this Gene-FC correlation was also obtained using unrelated participants from HCP S1200 release (Supplementary Sheet 17): *r* = 0.49, *p*_SA_ = 0.010, and another independent HCP-Aging neuroimaging dataset (Supplementary Sheet 18): *r* = 0.45, *p*_SA_ = 0.013.

**Fig. 2.**
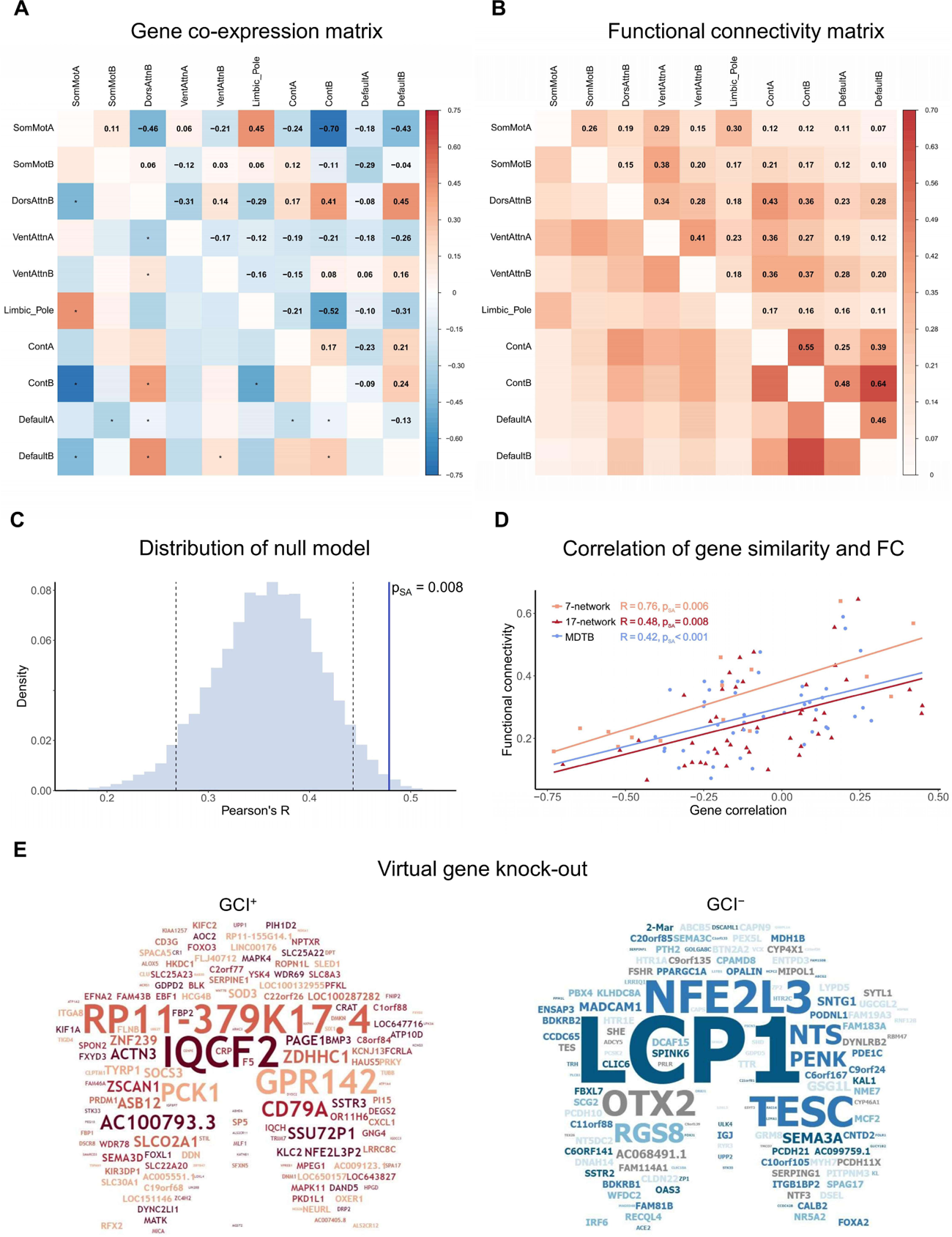
Network-specific gene co-expression correlates with functional connectivity (FC) within the cerebellum. **A** Genetic correlation was shown by the co-expression matrix (Supplementary Sheet 11) constructed for two bi-hemisphere donors across ten cerebellar networks using 443 cerebellar network-specific genes derived from all six donors. The 10 cerebellar networks corresponded to the networks containing samples from both bi-hemisphere donors (Table S3). \**p* ≤ 0.05 (Bonferroni corrected). **B** The FC matrix (Supplementary Sheet 12) shows the functional correlation for the ten cerebellar networks using 1,018 subjects from the HCP S1200 release [41]. All passed the significant threshold *p* ≤ 0.01 (Bonferroni corrected). **C** The probability distribution of null models preserving genetic spatial autocorrelation generated by BrainSMASH. The vertical black dashed lines correspond to the *p* values of 0.05 and 0.95; the blue vertical line showed the practical observed Gene-FC correlation (*r* = 0.48) and the corresponding *p*_SA_ = 0.008. **D** The overall intra-cerebellar Gene-FC correlation using different atlases: task-free 7-network (orange), 17-network (red) parcellation of the cerebellar functional atlas based on the cerebello-cortical FC, and task-based MDTB functional parcellation (blue) based on the task activation pattern. The Pearson’s correlation *r* and *p*_SA_ values are shown by the corresponding colors. **E** The GCI^+^ (*n* = 246, left, red) and GCI^−^ (*n* = 197, right, blue) gene lists were displayed on the flattened shape of the cerebellum. The word size and color intensity both indicate the GCI value.

Therefore, the 443 cerebellar network-specific genes that we derived based on the functional segregation of the cerebellum also correlated with the functional integration of the cerebellum. This Gene-FC correlation was not generated by the genetic SA, so it was consistently significant whenever using unrelated participants from HCP, independent HCP-Aging dataset, different parcellation resolution, or independent cerebellar functional atlas for the calculation. Moreover, the control test exhibited no Gene-FC correlation when the gene co-expression was constructed using non-network-specific genes (Supplementary Sheet 19) regardless of whether using a threshold or not. These findings confirmed that these 443 network-specific genes play a crucial role in intra-cerebellar functional organization.

### Convergently expressed genes among the cerebellar and cortical network-specific genes correlated with the FC across the cerebello-cortical cognitive-limbic networks

Since 90 of the 443 cerebellar network-specific genes were convergently expressed across the cerebello-cortical circuit, we wanted to know whether these ∼20% genes correlated with the FC across the cerebello-cortical circuit. A correspondence between the genetic and functional correlations was identified for the limbic (Fig. 3A): *r* = 0.36, *p* = 0.030 (FDR corrected), and control networks (Fig. 3B): *r* = −0.33, *p* = 0.034 (FDR corrected), but was not significant for the somatomotor: *r* = −0.15, *p* = 0.394 (FDR corrected), dorsal attention: *r* = −0.19, *p* = 0.281 (FDR corrected), ventral attention: *r* = −0.04, *p* = 0.779 (FDR corrected), or default: *r* = 0.10, *p* = 0.544 (FDR corrected) networks. The high cortical genetic similarity between the limbic system and the adjacent control network: *r* = −0.90, *p* < 0.001 (FDR corrected), somatomotor network: *r* = −0.55, *p* < 0.001 (FDR corrected), and ventral attention network: *r* = −0.72, *p* < 0.001 (FDR corrected) indicates that the gene co-expression between the cerebellar limbic network and the cortex reflects a gradual genetic gradient rather than genetic dissimilarity between the cerebellar limbic network and the other cerebellar networks. In addition, while controlling the effect of the cortical genetic similarity between the limbic and control networks, the partial correlation showed no cortical Gene-FC correlation for the control network: *r* = −0.13, *p* = 0.316. It implies that the significant cortical Gene-FC correlation for the control network could be explained by the high cortical genetic similarity between the cerebellar limbic and control networks. This observation is also consistent with the finding that convergently expressed genes were only observed in the limbic network but not in the control network (Table 1). Overall, these 443 cerebellar network-specific genes not only correlated with the intra-cerebellar FC, but ∼20% of them were also linked with the cerebello-cortical cognitive-limbic networks.

**Fig. 3.**
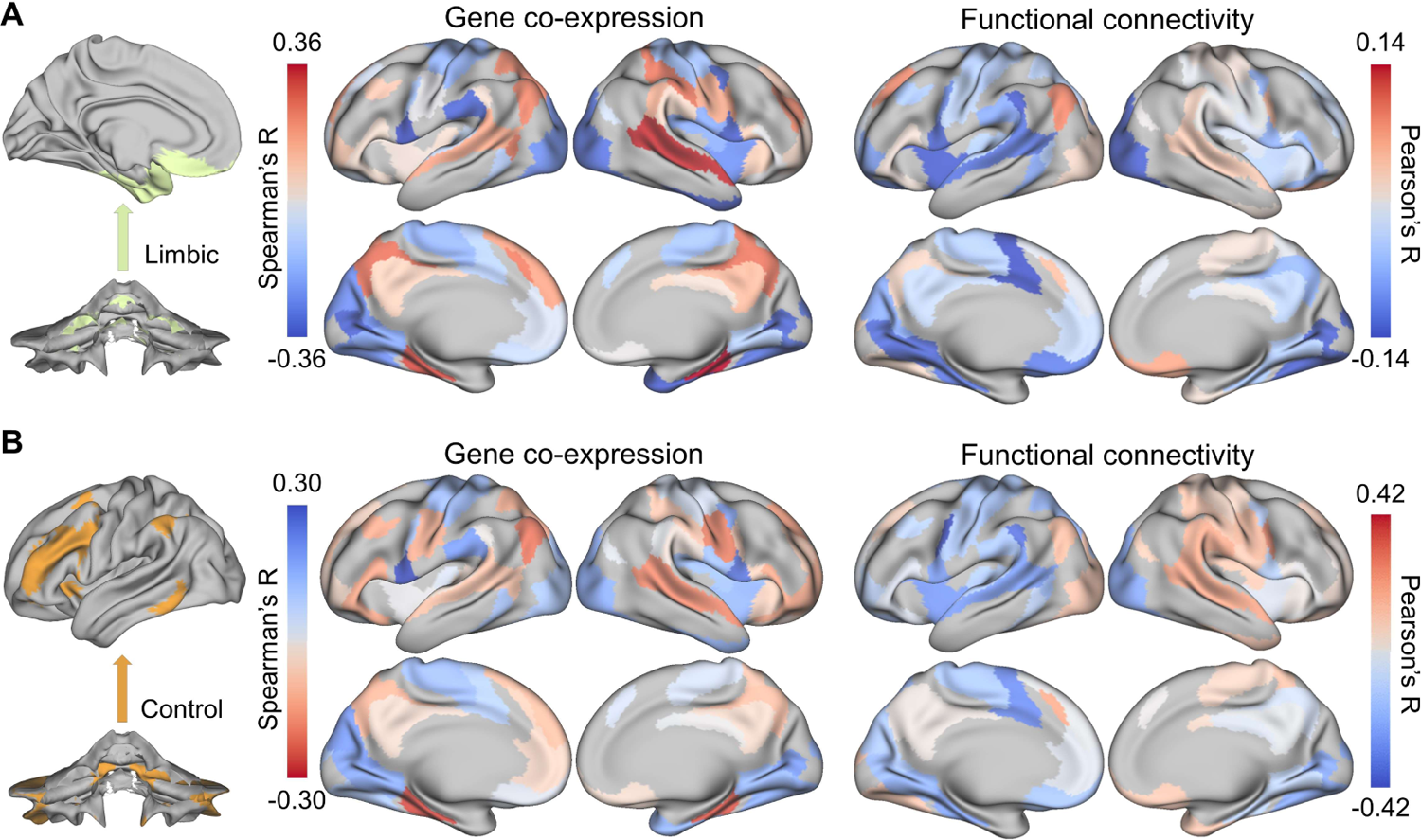
Cortical genetic and functional correlation of cerebellar limbic and control networks seeds. Both were calculated for two bi-hemisphere donors across six cerebellar networks and 59 cortical parcels that contained samples from both bi-hemisphere donors. **A** Limbic: the cortical gene co-expression (Supplementary Sheet 20) was calculated using the 90 overlapping genes between the cerebellar and cortical network-specific genes by Spearman’s correlation. The FC across each cerebellar network with each cortical parcel was calculated using Pearson’s correlation (Supplementary Sheet 21). The cortical limbic genetic and functional correlations were correlated with each other: *r* = 0.36, *p* = 0.030 (FDR corrected). **B** Control: the cortical gene co-expression and the FC for the control network were correlated with each other: r = −0.33, p = 0.034 (FDR corrected). Notice that the color bar of gene co-expression was inverted considering the negative Gene-FC correlation for the control network.

### Functional annotation revealed distinct biological properties of GCI^+^ and GCI^−^ separated by virtual KO

In addition to the overall correlation between gene co-expression and the functional integration of the human cerebellum, we investigated each gene’s importance to the intra-cerebellar Gene-FC correlation by scoring the 443 cerebellar network-specific genes based on the GCI. Using the virtual gene KO procedure (see more details in “Methods”), we were able to classify the 443 network-specific genes that linked cerebellar functional segregation and integration into two groups: a 246 GCI positive gene set (GCI^+^, Fig. 2E left and Supplementary Sheet 22) and a 197 GCI negative gene set (GCI^−^, Fig. 2E right and Supplementary Sheet 23). The distinction between the two sets is that the virtual KO of GCI^+^ genes increased the Gene-FC correlation, whereas the virtual KO of GCI^−^ genes decreased the Gene-FC correlation. Based on the winner-take-all principle, GCI^−^ genes may have a critical impact on the functional organization of the cerebellum. An example is that the top genes, LCP1 and TESC, enable GTPase binding and calcium binding, respectively [53], which are key functions within signaling transduction and consequently brain functions. Therefore, we applied a range of functional annotation tools to further explore the underlying roles of the GCI^+^ and GCI^−^.

#### Behavior-FC-Gene mapping analysis

Since these genes largely engaged in the cerebellar functional network which could predict human behaviors, whether they also involved in human behaviors with the FC acting as an intermediary role? We leveraged the Behavior-FC-Gene mapping analysis (see more details in methods) to explore the relationship between the co-expression of GCI^+^ and GCI^−^ with the human behaviors. Based on the PCA of 59 behavior measures (Table S6) across 211 HCP unrelated participants, 8 significant PCs were derived (Supplementary Sheet 24). Then the Behavior-FC map of each significant PC calculated by mass univariate regression procedure was compared with the co-expression pattern of GCI^+^ and GCI^−^ respectively (Supplementary Sheet 25). Among the eight significant PCs, we observed correlations of the Behavior-FC maps of PC1, PC3, and PC4 (Supplementary Sheet 25) with the GCI^−^ co-expression, rather than the GCI^+^.

But the PC1 and PC4 had few regression coefficients (1.47% and 0.74%, respectively) in the Behavior-FC map that survived the permutation test (Supplementary Sheet 25) compared with PC3 (Figs. 4B and S4). These significant Behavior-FC regression coefficients mainly occurred in the connections between default, control, and somatomotor networks with others (Fig. 4C), such as the connection between default B network with somatomotor A, somatomotor B, ventral attention A, limbic pole, and default A networks. Taking the intra-cerebellar FC as an intermediate role, PC3 had a significant correlation with the GCI^−^ co-expression (Fig. 4E, F): *r* = 0.67, permutation test *p* < 0.001. In contrast, no correlation between GCI^+^ co-expression and PC3 Behavior-FC map: *r* = 0.27, *p* > 0.05. Based on the loading pattern of PC3 for the original 59 behavior measures, the PC3 reflects the variation in the emotion-cognitional behavior measures (Fig. 4A). Specifically, PC3 has a large positive loading for the number of correct responses in emotion recognition, executive function, and fluid ability of cognition tasks, whereas it has a large negative loading for the response time in emotion recognition and fluid intelligence tasks. It suggests a high PC3 score might reflect high emotion and cognition ability. This Behavior-FC-Gene mapping analysis revealed that the GCI^−^ not only plays an important profiling role in the functional organization of the cerebellum but also implicates emotion-cognitional behavior vicariously through the intra-cerebellar FC.

**Fig. 4.**
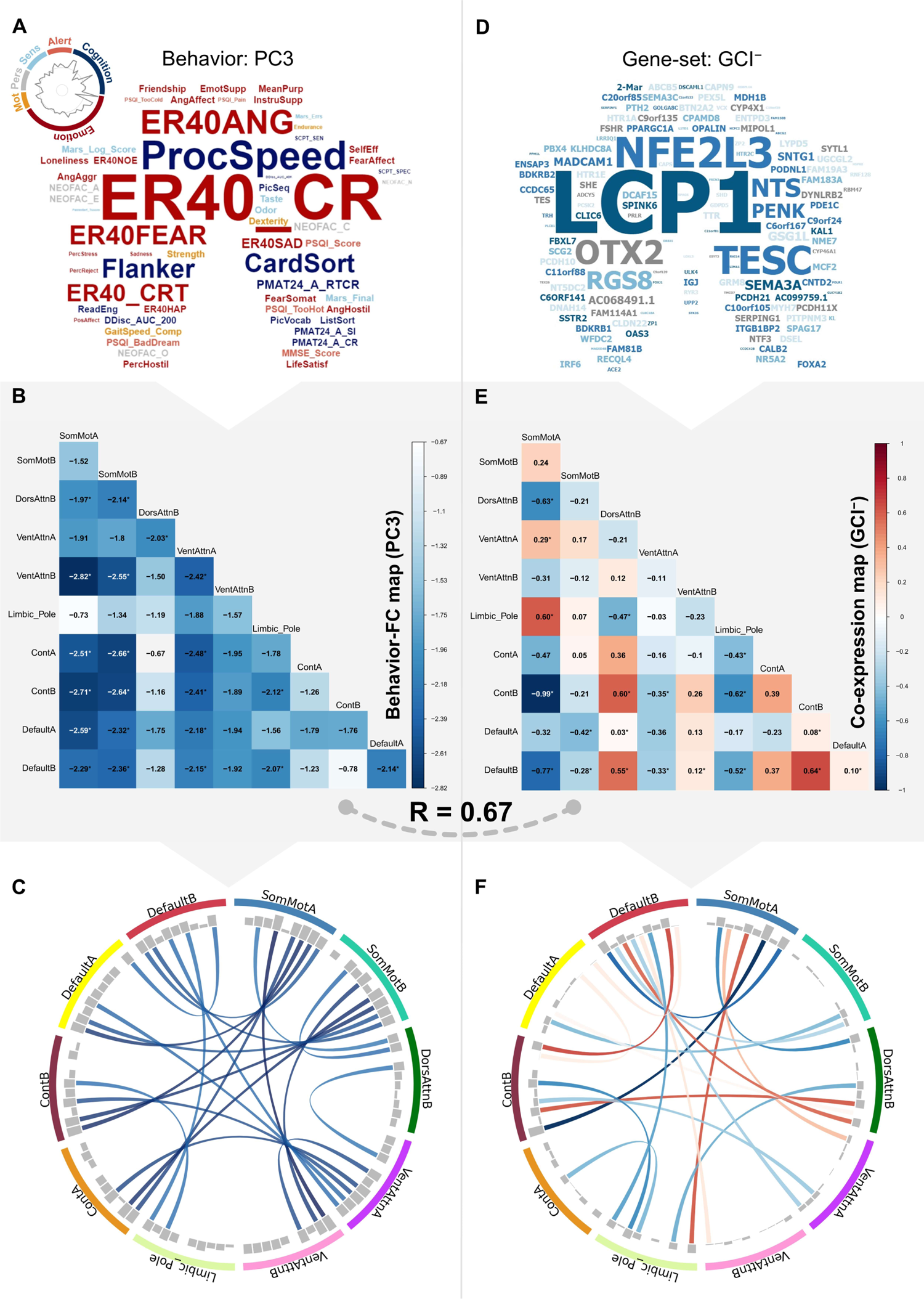
The Behavior-FC-Gene mapping analysis. **A** The loading pattern of principal component 3 (PC3) for the original 59 behavior measures (Table S6) was obtained from principal component analysis (PCA). The word sizes represent the absolute values of the loading coefficients, and the different colors indicate the six categories of the behaviors (Table S6). **B** Behavior-FC map for PC3, which are the z-scored regression coefficients of PC3 in each intra-cerebellar functional connection via mass univariate regression procedure. **C** The chord plot shows the significant regression coefficients after the permutation test consisting of 10,000 permutations. The circularly arranged gray histogram shows the absolute magnitudes of the regression coefficients, which correspond to the values of the regression coefficients shown in **B**. The color intensity of chords is the same as the color bar of **B**. **D** The GCI^−^ gene set which used to construct the GCI^−^ co-expression map **E**. **F** The chord plot shows the significant correlations: p ≤ 0.05 (Bonferroni corrected). The circularly arranged grey histogram shows the absolute magnitudes of the correlation coefficients, which correspond to the values of the correlation coefficients shown in **E**. The color intensity of chords is the same as the color bar of **E**.

### GO, pathway, and disorder enrichment analysis

The GO enrichment analysis of the GCI^+^ and GCI^−^ is shown in Fig. 5A. The GCI^+^ was mainly enriched in microtubule-related terms, including the microtubule associated complex (ID: 0005875), motile cilium (ID: 0031514), and dynein complex (ID: 0030286). Compared with GCI^+^, the GCI^−^ was not only enriched in microtubule-related terms but was also significantly enriched in terms related to neurotransmitter transport, such as calcium ion binding (ID: 0005509), regulation of hormone levels (ID: 0010817), response to catecholamine (ID: 0071869), response to monoamine (ID: 0071867), and regulation of neurotransmitter receptor activity (ID: 0099601). These findings are consistent with their different pathway enrichment results (Supplementary Sheets 26 and 27) in that the GCI^+^ was primarily enriched in some basic biological pathways: proximal tubule bicarbonate reclamation (ID: M4361) and glycolysis/gluconeogenesis (ID: M39474), which provides the energy needed during microtube-related processes. In contrast, the GCI^−^ was primarily involved in signaling transduction, especially in some neurotransmission pathways, such as the neuroactive ligand-receptor interaction (ID: M13380).

**Fig. 5.**
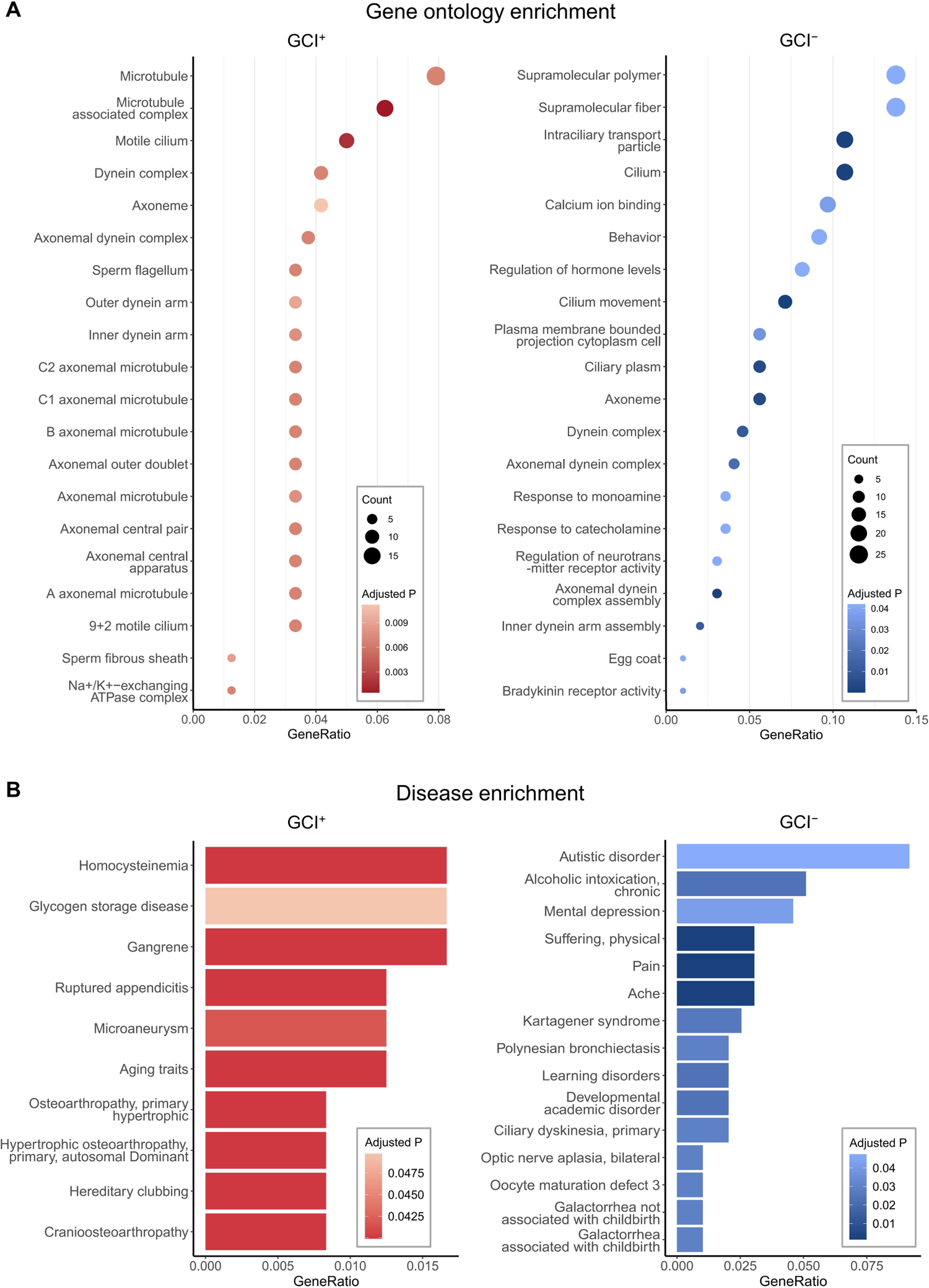
The gene ontology (GO) and disease enrichment analysis for GCI^+^ and GCI^−^. **A** Bubble plots showing the GO enrichment top 20 terms for GCI^+^ (left) and GCI^−^ (right) (Complete results are shown in Supplementary Sheets 26 and 27, respectively). The biological process (BP), cellular component (CC), and molecular function (MF) are displayed together. The dot size (count) represents the number of genes that are within the interest GCI^+^ or GCI^−^ gene panels as well as a specific GO term (*y*-axis). The different color intensities indicate the *p* value (FDR corrected). **B** Gradient bar plots showing the disease enrichment for all representative results for GCI^+^ and top 15 representative terms for GCI^−^. The different color intensities indicate the *p* value (FDR corrected).

Since the GCI^+^ and GCI^−^ are involved in different biological processes, we hypothesized that they also play different roles in brain disease or are related to different brain diseases. Unexpectedly, we found no link between GCI^+^ and any brain-related illnesses (Fig. 5B, left) but observed involvement of GCI^−^ in various neuropsychiatric disorders (Fig. 5B, right), including autistic disorder (ID: C0004325), alcoholic intoxication (ID: C0001973), mental depression (ID: C0011570), pain (ID: C0030193), learning disorders (ID: C0023186) and others (Supplementary Sheet 27). Many of these, especially mental depression and autistic disorder, have a close relationship with the human cerebellum, in which patients have shown FC abnormalities [54, 55]. The mental depression- and autistic disorder-associated genes were TRH, PENK, TTR, ADCY5, NRXN1, HTR1A, HTR2C, NTS, PEX5L (*n* = 9, Supplementary Sheet 27), and DLGAP2, TRH, PENK, RYR3, SEMA3A, NRXN1, TESC, ABCG2, PCDH10, CNTN4, HTR1A, CALB2, HTR2C, DNAAF4, FOLR1, NTS, GRM8, UPP2 (*n* = 18, Supplementary Sheet 27), respectively, and the overlapping genes were TRH, PENK, NRXN1, HTR1A, HTR2C, NTS (*n* = 6).

### Integrative temporal specificity analysis

In light of the distinct properties of GCI^+^ and GCI^−^, we wanted to know whether the roles played by these two gene sets showed variable prevalence at different ages. By leveraging the BrainSpan dataset [56] and applying the analysis strategy of the CSEA tool [52], we found that GCI^+^ showed significant overexpression in early middle fetal, late middle fetal, late fetal, neonatal early infancy, and adolescence compared with GCI^−^ (Fig. 6A). These stages neatly correspond to the timeline of the protracted development of the human cerebellum [57], which extends from the early embryonic period until the end of the first postnatal year. This result appears to be consistent with the observation that the GCI^+^ is involved in some fundamental biological processes, especially microtubule-related activity, whose dynamics play a key role in cerebellar neurodevelopment [58]. In contrast, compared with the GCI^+^, the GCI^−^ was significantly expressed in early mid fetal, neonatal early infancy, late infancy, early childhood, adolescence, and young adulthood (Fig. 6B). These periods include the highest neurodevelopmental risk windows for autism spectrum disorder (ASD) [59] and major depression disorder (MDD) [60], both of which also existed in the disease enrichment analysis of GCI^−^.

**Fig. 6.**
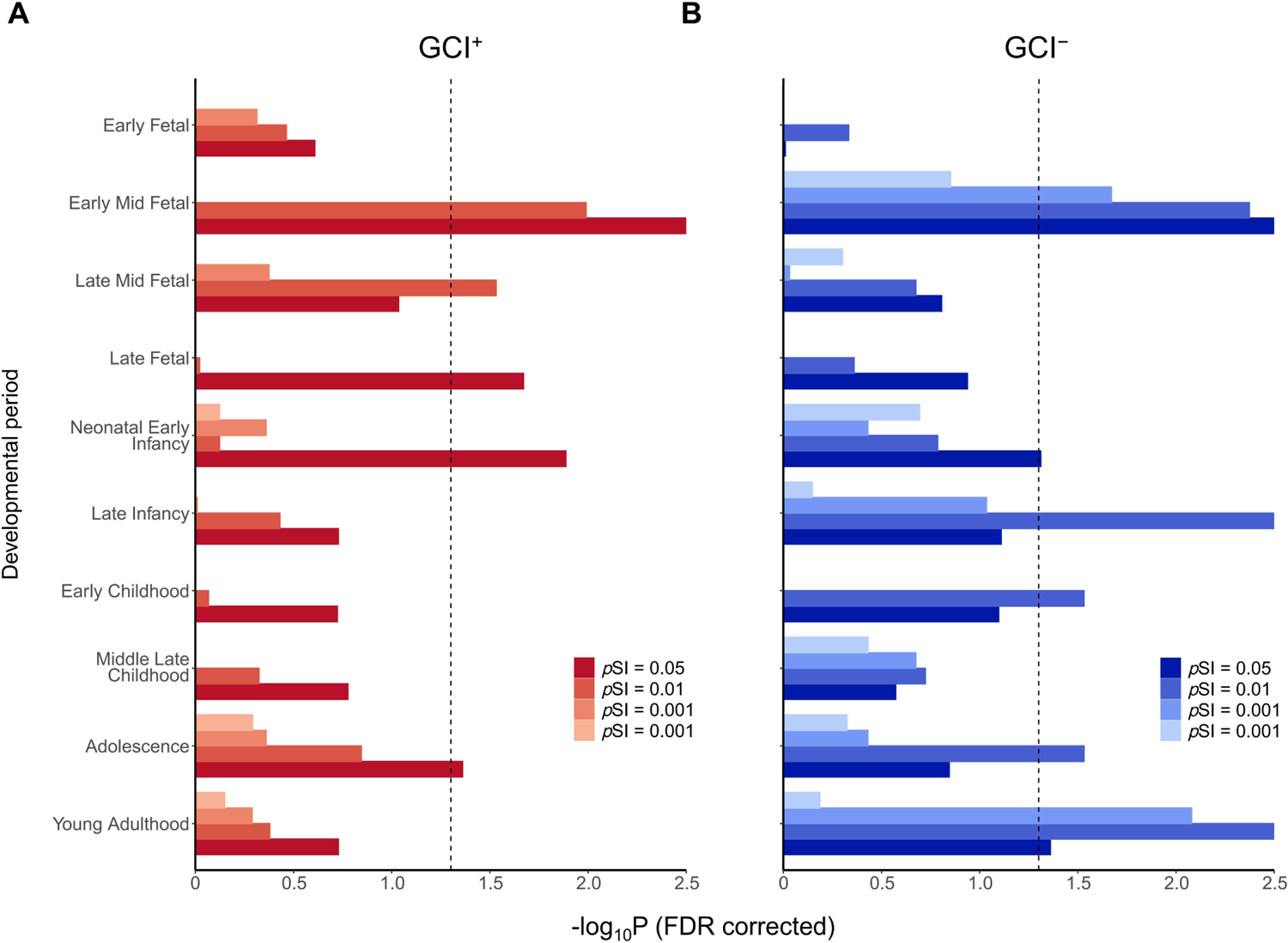
Integrative temporal specificity analysis within the human cerebellum for GCI^+^ and GCI^−^. **A** Bar plot showing the temporal specificity results for GCI^+^. The specificity index probability (pSI = 0.05, 0.01, 0.001, and 0.0001, permutation corrected, shown as different color intensities) was used to determine how likely a gene was to be expressed in a given time window relative to all other time windows [52]. The *x*-axis corresponds to the -log10*p* (FDR corrected), and for esthetics if -log10*p* (FDR corrected) > 2.5, -log10*p* (FDR corrected) = 2.5; the y-axis represents the 10 development windows collected by BrainSpan [56]. The vertical dark dashed line corresponds to the *p* (FDR corrected) = 0.05. Complete results are shown in Supplementary Sheet 28. **B** Bar plot showing the temporal specificity results for GCI^−^.

## Discussion

The current study provided a tentative exploration of the genetic differential and co-expression linked with the functional organization of the human cerebellum and has the potential for elaborating and rethinking the neurobiological underpinnings of the cerebellar functional organization. Furthermore, we identified two gene sets involved in cerebellar neurodevelopment and neurotransmission and found indirect but interesting genetic evidence supporting a key role played by the cerebellar functional network in emotion-cognitional behaviors, as well as many neuropsychiatric disorders. These findings hint at a micro-macro interacted mechanistic possibility for the cerebellar contributions to emotion-cognitional behaviors, and consequently to related neuropsychiatric disorders.

### The genetic profiles underlying cerebellar functional segregation correlate with intra-cerebellar and cerebello-cerebral connections

In this study, we found correlations between the identified cerebellar network-specific genes with the intra-cerebellar connection and cerebello-cerebral FC. These findings could provide possible empirical genetic support for the hypothesized decisive role of cerebellar connectivity in the functional heterogeneity of the cerebellum. First, while obtaining the network-specific genes, we found significant differences in the number of identified genes between the functional specificity (i.e., limbic, visual networks) and functional diversity networks (i.e., the control, default networks); specifically, more differentially expressed genes were in the former and vice versa in the latter [61]. This observation was also found in a previous cortical gene expression homogeneity analysis [9] that showed that a relatively high differential expression pattern was observed in the primary sensory cortex, area 38, and the primary visual cortex. But the findings related to the inconsistency in the amount of somatomotor cerebellar (*n* = 3) and somatomotor cortical network-specific genes (*n* = 960) were unclear. One possible explanation may be that the preferential links between the cerebellar representations of body space and the motor, somatosensory, and premotor cortices are challenging to distinguish [21]. The finding of cerebellar network-specific genes is consistent with the elaborate regional difference in cerebellar-cortical cytoarchitecture [2, 62] in addition to the uniform cell types and their connectivity, such as regional variation in the cell size and packing density of Purkinje, granule, and Golgi cells. Second, the overall distribution patterns of the cerebellar and cortical network-specific genes were highly correlated, a finding in line with a similar macroscale principle identified in the cerebellar and cortical functional organization [23, 63]. These correlated patterns may be related to defining the cerebellar networks, which were by projecting the cerebral cortical networks onto the cerebellum by computing the functional connections between the two structures [21]. More interestingly, the molecular genetic substrates simultaneously linking functional heterogeneity and integration could be observed across different functional subdivisions, regardless of whether the parcellation was based on the task-free cerebello-cortical FC [21] or the intra-cerebellar task-based activation pattern [20]. These interpretations are further supported by the widely accepted notion about the human cerebellum that its functional specialization is dominated more by its connection with extra-cerebellar structures than within its generally homogeneous cytoarchitecture [8]. Although no intra-cerebellar anatomical fiber connections linking adjacent or distant cerebellar regions have been observed [64, 65], it is well recognized that the intra-cerebellar functional map is a consequence of the topological arrangement of its extra-cerebellar anatomical connections [8]. This fact is especially important to note when explaining the Behavior-FC results that although our results were obtained through intra-cerebellar connections, the cerebellar-cortical connections must also be involved actually. The proposed relationship between extra- and intra-cerebellar connectivity, in turn, can be expected to affect the resting-state activity between cerebellar regions [23]. Furthermore, for the interaction between gene and FC, they influence one another bidirectional [66] through gene-environment (G-E) interplay [67] with the development as an essential element [68], instead of a unidirectional determination role of the genes in the brain connection. Here we observed the Gene-FC interaction from the adult perspective (AHBA donors aged from 24 to 57 years old), which lacks exploration in the G-E interplay compared with the early development [68].

Third, in addition to the intra-cerebellar Gene-FC correlation, we observed a direct correlation between genes underlying the cerebellar functional specialization and cerebello-cerebral FC of the limbic and control networks. The Gene-FC correlation in the control network was mainly caused by the genetic similarity between these two networks; this interaction between limbic-emotion and control-cognition has been confirmed anatomically and behaviorally [69]. For instance, the integrated processing by the emotion and cognition areas has been identified solely based on their anatomical connections [70]. This relationship can also be observed in that when looking at the top of a hill, a sad mood induces a steeper perception of the hill than a happy one [71]. One possible reason why we only obtained this correspondence in the limbic network may be the low functional heterogeneity [61] and inter-individual functional variability [72] of the limbic network compared with others as well as the complexity of gene expression; i.e., the Gene-FC correlation is not fully portrayed by the differentially expressed genes [26]. Considering the indirect connection between cerebellum and cortex, and large differences between the cerebellum and cortex in terms of their gene expression patterns [13] and structure-function relationships [73], as well as the individual variability of their functional networks [75], the convergently expressed genes associated with the cerebello-cortical cognitive-limbic networks were evident through simple analysis of co-expression, demonstrating these genes are very significant and may hold clues to the molecular underpinnings of the cognitive-emotion roles played by the cerebello-cortical circuit. For example, the HTR1A and HTR2C, both preferentially expressed in the cerebellar and cortical limbic network, are pivotal genes in serotonin transmission and play a modulation role in the limbic system act as important therapeutic targets in limbic system-related disorders [74]. This observation was further confirmed by the intra-cerebellar Behavior-FC-Gene mapping analysis, which showed that some of the genes were strongly associated with the emotion-cognitional behaviors.

### Cerebellar neurodevelopment feature of GCI^+^, cerebellar neurotransmission, emotion-cognitional behavior, and neuropsychiatric related features of GCI^−^

Interestingly, we identified two gene subsets with pronouncedly different characteristics based solely on the direction in which each gene influenced the intra-cerebellar Gene-FC correlation by applying a simple virtual KO approach on the 443 cerebellar network-specific genes. By using a series of bioinformatic tools, we found converging evidence for GCI^+^ and GCI^−^ involvement in cerebellar neurodevelopment and cerebellar neurotransmission, respectively. It is also interesting to speculate that these 443 network-specific genes that link both cerebellar functional segregation and integration have a relationship with some brain disorders since prior evidence showed that the cerebellar functional organization plays a key role in various neurological [30, 75] and psychiatric disorders [29], most of which possess common underlying genetic risks [76]. But a tricky problem emerged in that the genes we are interested in were derived from healthy individuals. It could be tackled to some extent by using the virtual KO method, which can simulate the different expression levels of each gene and thus coarsely corresponds to a fraction of the expression level under normal health and disease situations. This is why we thought that we might be able to see whether the GCI^+^ and GCI^−^ are related to a specific disease even though the genes were derived from healthy individuals.

The GCI^+^ is involved in many microtubule-related terms and is overexpressed throughout the protracted development of the cerebellum. The dynamics and flexibility of microtubules were found to be essential throughout cerebellar development via leading the morphological alterations of Purkinje cells [58]. In addition, some genes of the GCI^+^, such as GTPBP2 [77] and Lin28b [78], were found to play a key role in neurodevelopment; overexpression of the Lin28b gene can induce the development of pathological lobulation in the cerebellum [78]. This converging evidence prompts our speculation that the GCI^+^ is engaged in cerebellar neurodevelopment. Unexpectedly, the GCI^+^ showed no link to brain-related diseases, which appears to be consistent with its primary involvement in many fundamental biological functions. However, this lack of disease linkage is inconsistent with the significant overexpression of GCI^+^ genes during the protracted development of the cerebellum. Many researchers pointed out that this protracted development increased the susceptibility of the cerebellum to many psychiatric disorders [57]. It is likely complemented by the overexpression of GCI^−^ in the early middle fetal and neonatal early infancy periods. Other possible explanations include few genetic studies of the cerebellum compared with the cerebral cortex, and large gene expression differences between the cerebellum and extra-cerebellar structures [13], so the related datasets may lack sufficient information specific to the cerebellum. It calls for future studies to provide a complete explanation by considering multiple perspectives.

The GCI^−^ was found to be related to emotion-cognitional behaviors, involved in many neurotransmission processes, enriched in various neurological and psychiatric disorders, and significantly overexpressed in late infancy, early childhood, adolescence, and young adulthood compared with GCI^+^. These results are mutually supportive. The neurotransmission has long been believed to play a crucial role in emotional [79] and cognitional behaviors [80], and its abnormality has been extensively associated with various neurological [81] and psychiatric disorders [82, 83] with the characteristic of cognitive and emotional impairments. For example, the abnormal transmission of monamines and catecholamines, such as serotonin and dopamine, has been widely linked with many psychiatric disorders, and these transmitters have thus become potential treatment targets [84]. The period through which the GCI^−^ genes are expressed includes the high-risk time windows for GCI^−^ enriched disorders, such as mental depression (aged 18–29) [60] and autistic disorder (from infancy to childhood) [59]. And the high expression of GCI^−^ in early middle fetal life might be associated with the prenatal risk factors associated with depression [85] and autism [86].

### A possible micro-macro interacted mechanistic explanation for the function/dysfunction of the human cerebellum

The cerebellum regulates motor, emotion, and cognition by functioning as an oscillation dampener that combines multiple internal representations with external stimuli and appropriate responses to maintain the behavior around a homeostatic baseline automatically to optimize performance according to the cortex [87]. Interestingly, we found that the GCI^−^ associated with the emotion-cognitional behaviors via the intra-cerebellar FC, mainly includes the connections between control, default, and somatomotor networks which have been widely confirmed to be involved in these behaviors via human clinical data or functional imaging studies [4, 88–90]. This result suggests that the interaction between genes and FC is also involved in various higher functions of the cerebellum, providing new evidence and molecular substrates for the progressively complete notion that the cerebellum modulates cognition and emotion as it does to the motor control [1]. Beyond that, the specifics of how the cerebellum is involved in emotional and cognitive functions are explained by many compatible theories [87]. Such as the internal model [91], in which the cerebellum shows increased activations for negative feedback compared to the positive feedback in the reversal-learning task [92], which is maybe consistent with the negative Behavior-FC map found in the present study.

Meanwhile, the regions that showed a significant correlation between emotion-cognitional behaviors and GCI^−^ via connections, correspond exactly to the cerebellar posterior lobe whose lesions produce the dysmetria of thought and emotion, i.e., the cerebellar cognitive affective syndrome, which is characterized by the deficits in executive function, visual-spatial processing, linguistic skills and affect regulation [93]. And no significant difference in the performance on newly established objective cognitive tests battery between patients with isolated cerebellar lesions versus complex cerebello-cortical connection disorders, such as schizophrenia, mental depression, and autistic disorder [94]. It is thus reasonable and mutually validating that we found that the GCI^−^ enriched in many neuropsychiatric disorders involving alternation in these two categories of behaviors, including mental depression, autistic disorder, pain, alcoholic intoxication, learning disorder, and others. Moreover, these disorders are all closely related to the alterations of the cerebellar FC [29, 54, 95–97].

Therefore, the GCI^−^ provides a possible micro-macro interacted mechanistic explanation for the functions and dysfunctions of the human cerebellum. The genes underlying the functional organization of the human cerebellum are likely involved in emotion-cognitional behaviors through their interaction with the cerebellar connection. One of the possible ways the risk genes act in the pathogenesis of corresponding diseases maybe through their interactions with the cerebellar FC, which results in dysregulation of cerebellar FC and thus pathologically manifested as its functional connection abnormalities and consequently transient or long-term impairment at the cognitive and emotional behavioral levels of these diseases. For instance, the fluctuation in the correspondence of the Gene-FC relationship found in the present study and widespread altered cerebellar FC excavated previously in diverse neuropsychiatric disorders [29]. The GCI^−^ also provides a promising genetic resource for investigating the cerebellar involvements in emotion-cognitional behaviors and related brain diseases. For example, the observed overlapping genes, i.e., NRXN1, are associated with mental depression and autistic disorder supported previous clinical studies showing that rare and common variants in NRXN1 carried risks for MDD [98], ASD, and schizophrenia [99]. And HTR1A, which has a high expression in the cerebellum [14] and has been linked to the cognitive process in both animal models and human [79], was found to be involved in pain, mental depression, autistic disorder, alcoholic intoxication, learning disorder, and other conditions (Supplementary Sheet 27).

### Methodological considerations

The interpretation of our findings is not without limitations. First, we validated our results across independent imaging datasets that cover the age range of AHBA, and various cerebellar atlases with different parcellation criteria and resolutions. But we are unable to test our results in an independent gene dataset as there is no other publicly available dataset with a detailed sampling of subregions of the human cerebellum. Second, the gene co-expression we constructed only considered one small part of the relationship between the genes and FC. Thereby it did not fully recapitulate the complexity of the brain transcriptome, such as gene-gene interactions [100]. That is one possible reason we only found a cerebello-cortical Gene-FC correlation for the cognitive-limbic networks. Third, the simple correlation approach [25, 61] used in this study and other linear regression models like partial least squares analysis [101], can only prioritize genes for further investigation and cannot fully explore the causal relationship between genes and functional organization. As a result, further exploration is hindered by the intricacies of genetic and epigenetic regulation. It makes the discussion and explanation of the different directions of this correlation challenging. For example, why the direction of influence on the Gene-FC correlation could separate these 443 genes into two distinct gene sets with different functions remains unclear. Hence further related exploration is necessary but very challenging. Fourth, since the 443 genes were not derived based on the correlation with a spatially-defined phenotype, the newly proposed strategy [102] that mitigates the bias of leveraging gene enrichment approach in the spatial transcriptomic data cannot be directly utilized. Nonetheless, it cannot be determined whether this type of bias affects the present results, which calls for future efforts to develop methods to answer this question.

Briefly, more donors, the overall pattern of gene expression, gene regulation, epigenomics, and improved cellular resolution are needed and imperative for developing more appropriate and ingenious approaches to understand the bidirectional relationship between genes and functional organization, which is a greater challenge for neuroscience than just identifying a link between genetic and imaging data. Nevertheless, in light of the currently limited understanding of how microscale genes contribute to macroscale brain functional organization, the prioritization of genes and the related functional annotations presented here are still necessary [24, 26].

## Conclusions

In summary, we found that the network-specific genes underlying cerebellar functional heterogeneity correlated with the intra-cerebellar and cerebello-cerebral FC. It indicates that the genetic infrastructure associated with functional segregation coalesces to form a collective system, which closely relates to the functional integration of these functional subregions. The current study has thus unveiled part of the neurobiological genetic substrates underlying the cerebellar functional organization. We also identified important indirect genetic markers that support the key role played by the cerebellar functional network in emotion-cognitional behaviors and many brain disorders. These findings hint at the possibility of establishing a cerebellar “gene—connection—function/dysfunction” chain, as well as of helping to bridge the knowledge gap between the genetic mechanisms driving the cerebellar functional organization and the predictive markers of behaviors and heritable risks of disorders, especially major depression and autistic disorder.

## Acknowledgements

This work was partially supported by the Science and Technology Innovation 2030—Brain Science and Brain-Inspired Intelligence Project (Grant No. 2021ZD0200203), the Natural Science Foundation of China (Grant Nos. 82072099 and 91432302), the Strategic Priority Research Program of the Chinese Academy of Sciences (XDB32030214), the Youth Innovation Promotion Association, the Beijing Advanced Discipline Fund and the Major Scientific Project of Zhejiang Lab (Grant No. 2021ND0PI01). Data were provided by the Human Connectome Project, WU-Minn Consortium (Principal Investigators: David Van Essen, and Kamil Ugurbil; 1U54MH091657) funded by the 16 NIH Institutes and Centers that support the NIH Blueprint for Neuroscience Research; and by the McDonnell Center for Systems Neuroscience at Washington University. The authors appreciate the English language and editing assistance of Rhoda E. and Edmund F. Perozzi, PhDs.

## Author Contributions

YW and LF designed the research; YW and LC performed the experiments; YW, LF, CC, TJ, BL, and ZY contributed new analytic tools; YW, LC, DL, XW and CG analyzed data; YW and LF wrote the paper; and LF, CC, KHM, JRN, YZ, ZY, JX, SBE, BL, and TJ contributed analytic expertise, theoretical guidance, paper revisions, and informed interpretation of the results; and LF and CC supervised the project.

