## Supplementary material for "Uncovering the Genetic Profiles Underlying the Intrinsic Organization of the Human Cerebellum": Supplenmentary information

**This PDF file includes:**

Figure S1 to S4

Table S1 to S6

Legends for Supplementary sheet 1 to sheet 28

**Other Supplementary Materials for this manuscript include:**

Supplementary sheet 1 to sheet 28

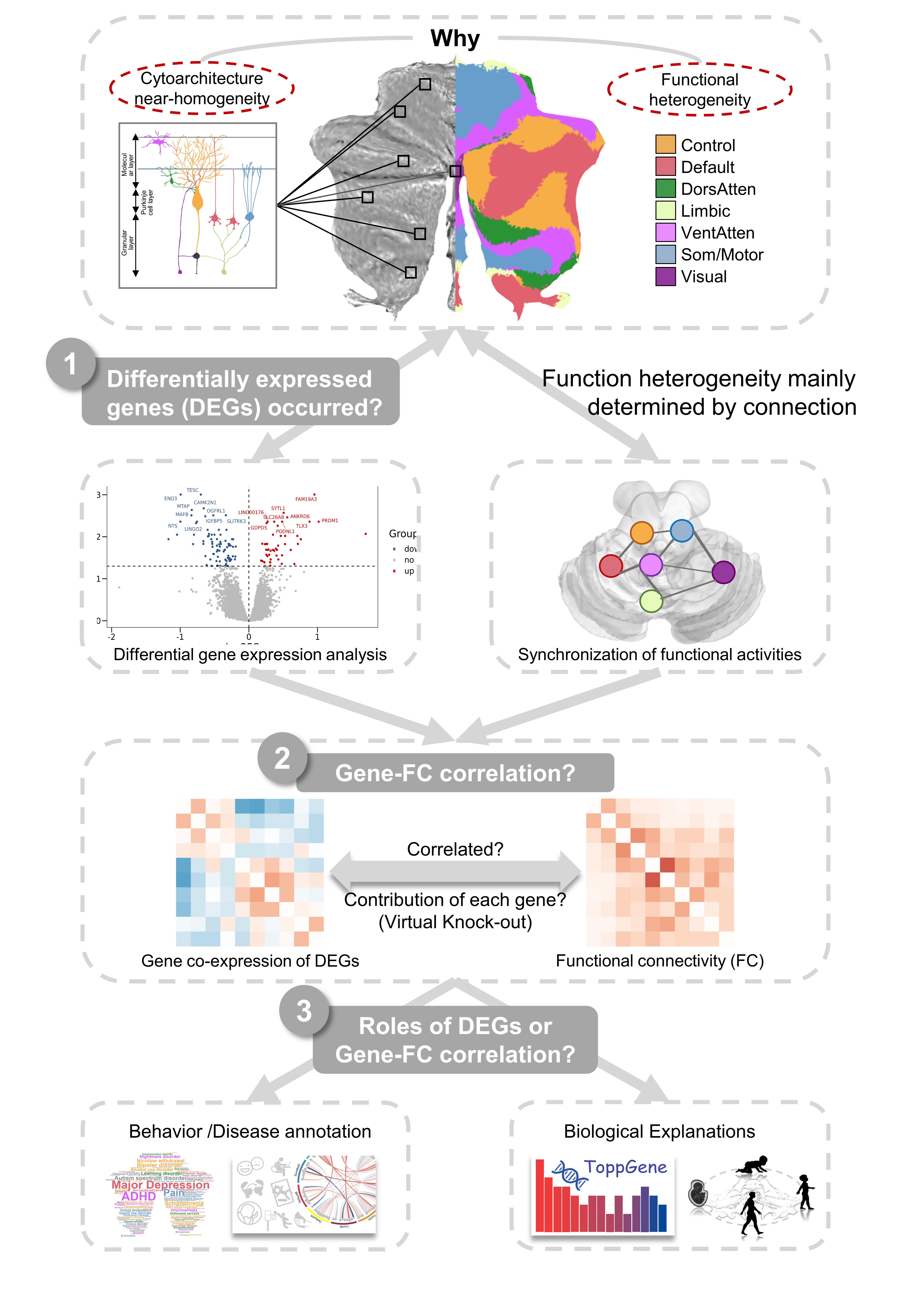

**Figure S1.** **Unraveling the genetic profiles underlying the cerebellar intrinsic organization by exploring three progressive questions** (related to Figure 1). Briefly, our scientific question is rooted in the inconsistency between cytoarchitecture near-homogeneity and functional heterogeneity of the human cerebellum. Since the macroscopic functional organization of the human brain is thought to be ultimately regulated by underlying microscopic gene expression [1], and the converging evidence has tended to support the view that the renewed functional diversity of the human cerebellum is derived primarily from its extensive connections with a preference for motor control, cognition, and emotion, rather than being limited to a uniform cerebellar cortical cytoarchitecture [2]. We hypothesized it could be explored from the analysis of differentially expressed genes (DEGs, Figure S1.1) and their relationship with functional connectivity (Gene-FC correlation, Figure S1.2). Lastly, the relationship between the DEGs or Gene-FC correlation with the human behaviors and disorders (Figure S1.3) could further link the genetic and behavior markers of the cerebellar functional network to its involvements in higher order non-motor functions and dysfunctions in related neuropsychiatric disorders.

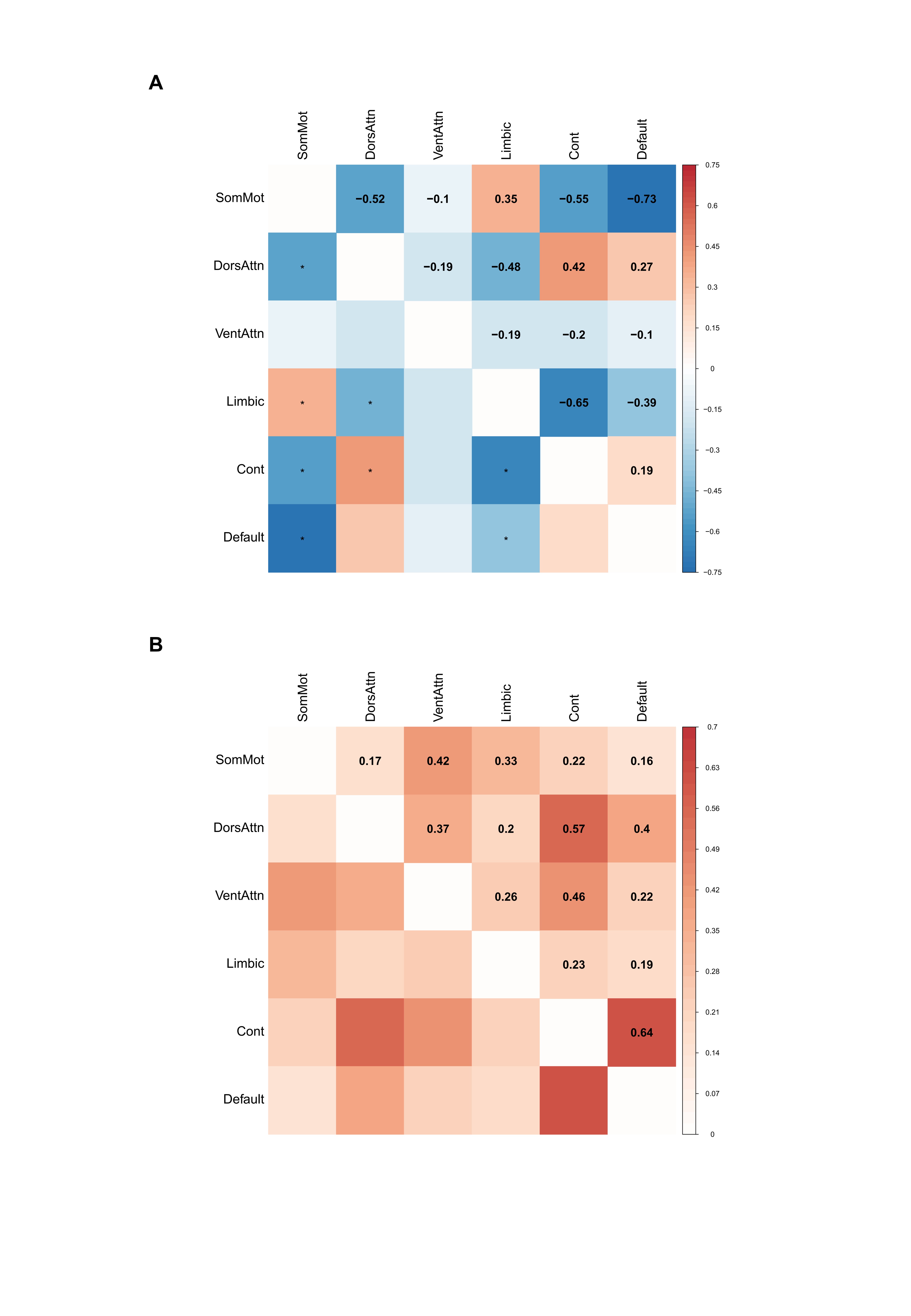

Figure S2. The intra-cerebellar Gene-FC correlation was validated using task-free 7-network parcellation [3] (related to Figure 2). (A) Genetic correlation is shown by the co-expression matrix (Supplementary sheet 13) constructed for the two both-hemisphere donors using 443 cerebellar network-specific genes derived from all six donors, and across 6 cerebellar networks which contained samples from both 2 bi-hemisphere donors. (B) FC matrix (Supplementary sheet 14) shows the functional correlation for the 6 cerebellum networks using 1,018 subjects from the HCP S1200 release (Van Essen et al., 2012). They correlated with each other, Pearson’s r = 0.76, p < .001.

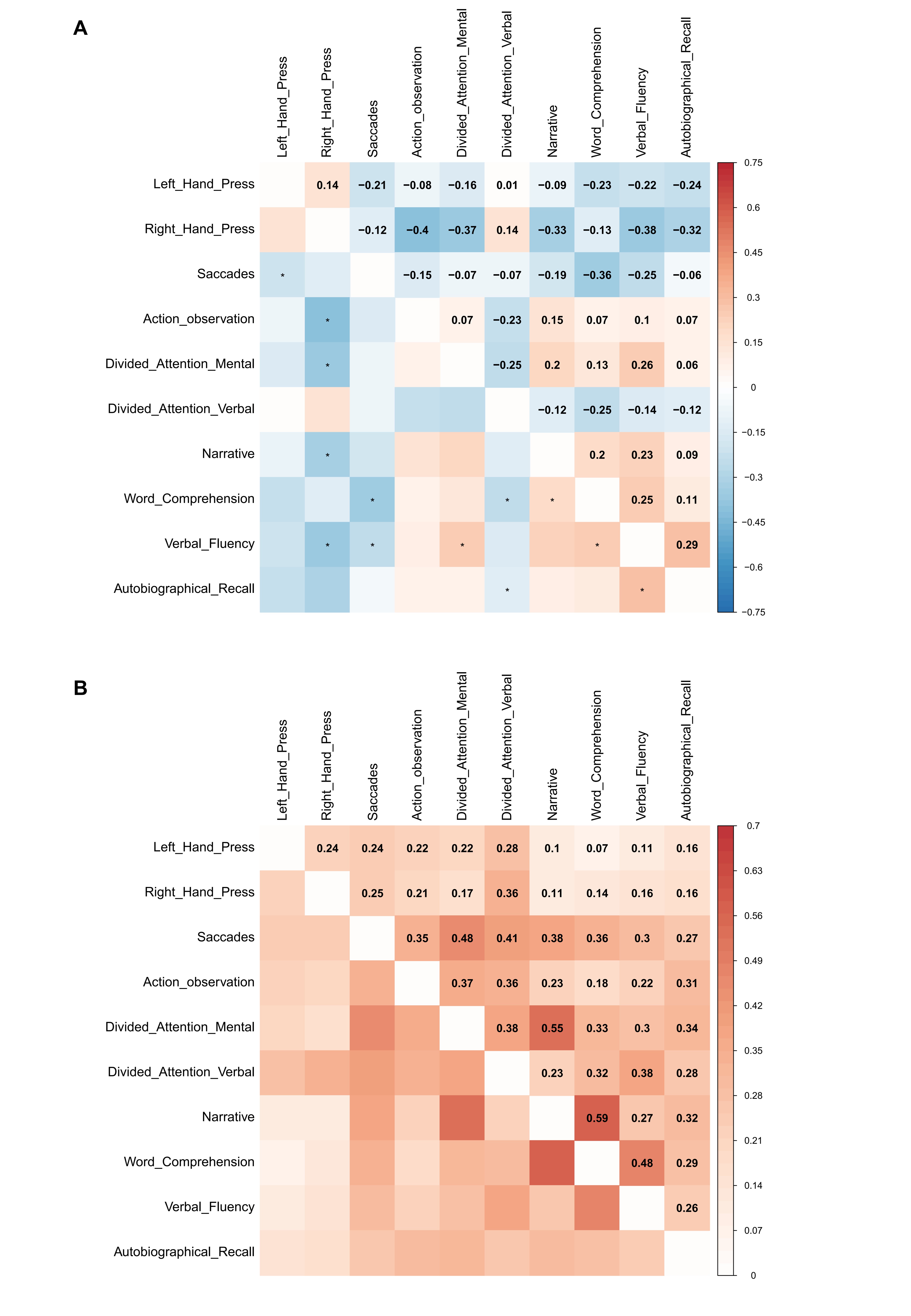

Figure S3. The intra-cerebellar Gene-FC correlation was validated using independent task-based Multi-Domain Task Battery (MDTB) functional parcellation, which is based on the task activation patterns [4] (related to Figure 2). (A) Genetic correlation is shown by the co-expression matrix (Supplementary sheet 15) constructed for the two both-hemisphere donors across 10 cerebellar regions using 481 cerebellar network-specific genes (Supplementary sheet 4) derived from all six donors. (B) FC matrix (Supplementary sheet 16) shows the functional correlation for the 10 cerebellum regions using 1,018 subjects from the HCP S1200 release (Van Essen et al., 2012). They correlated with each other, Pearson’s r = 0.42, p = .004.

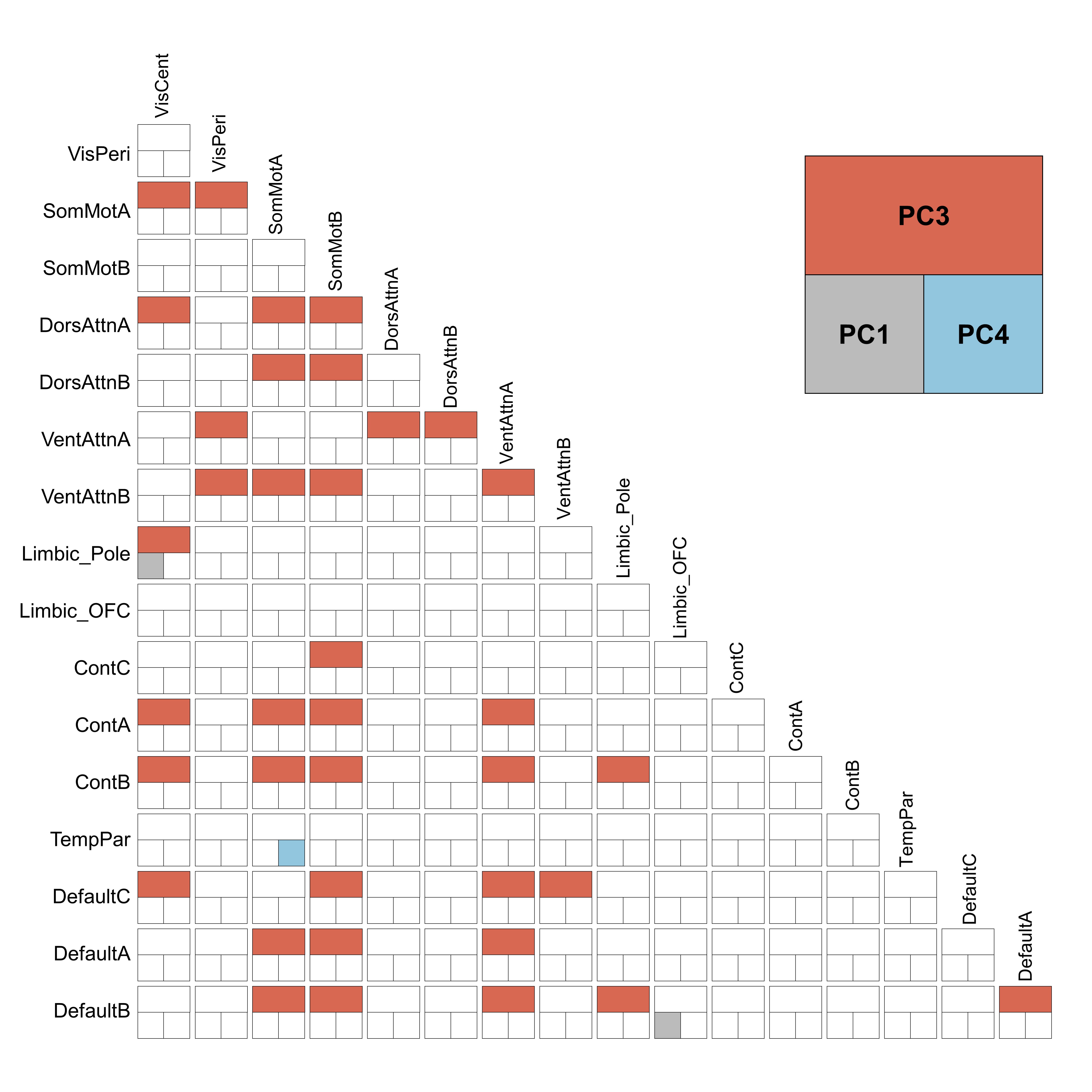

**Figure S4. The PC3 has a more significant correlation with the intra-cerebellar FC compared with the PC1 and PC4** (related to Figure 4)**.** The relationship between significant behavior PCs and each intra-cerebellar FC was investigated using a mass univariate regression procedure [5] across 211 unrelated subjects. The significance of the derived regression coefficients was assessed by a permutation test consisting of 10,000 permutations which randomly shuffled the order of the subjects for each significant PC score. The significant correlation between each intra-cerebellar FC and PC1 (grey), PC3 (orange), and PC4 (blue) were shown in the triangle of this 17 × 17 network matrix.

Table S1. Demographic information for the AHBA donors [1].

| Donors ID | Age | Ethnicity | Sex | L/R hemisphere |
| --- | --- | --- | --- | --- |
| Donor 9861 | 24 | Black/African American | M | L/R |
| Donor 10021 | 39 | Black/African American | M | L/R |
| Donor 12876 | 57 | White/Caucasian | M | L |
| Donor 14380 | 31 | White/Caucasian | M | L |
| Donor 15496 | 49 | Hispanic | F | L |
| Donor 15697 | 55 | White/Caucasian | M | L |

Donors 10021 and 9861 had bi-hemispheric data; the other 4 donors only had left-hemisphere data.

Table S2. Counts of the cerebellar samples falling within each functional network of the cerebellar 7-network population atlas [3].

| Donors ID  Network names | | 9861 | 10021 | 12876 | 14380 | 15496 | 15697 |
| --- | --- | --- | --- | --- | --- | --- | --- |
| Control |  | 9 | 15 | 7 | 5 | 8 | 10 |
| Default |  | 6 | 16 | 8 | 13 | 10 | 9 |
| Dorsal Attention |  | 3 | 7 | 8 | 8 | 5 | 7 |
| Limbic |  | 7 | 8 | 4 | 4 | 10 | 6 |
| Ventral Attention |  | 7 | 16 | 3 | 7 | 14 | 16 |
| SomatoMotor |  | 9 | 10 | 0 | 4 | 9 | 16 |
| Visual |  | 0 | 2 | 4 | 3 | 1 | 1 |
| None |  | 0 | 2 | 6 | 0 | 3 | 11 |

Table S3. Counts of the cerebellar samples falling within each functional network of the cerebellar 17-network population atlas [3].

| Donors ID  Network names | | 9861 | 10021 | 12876 | 14380 | 15496 | 15697 |
| --- | --- | --- | --- | --- | --- | --- | --- |
| ContA |  | 3 | 6 | 2 | 2 | 4 | 7 |
| ContB |  | 8 | 9 | 5 | 7 | 5 | 8 |
| ContC |  | 0 | 1 | 1 | 0 | 1 | 0 |
| DefaultA |  | 2 | 9 | 4 | 4 | 5 | 0 |
| DefaultB |  | 2 | 6 | 1 | 5 | 4 | 7 |
| DefaultC |  | 0 | 0 | 3 | 4 | 3 | 0 |
| DorsAttnA |  | 1 | 0 | 0 | 0 | 1 | 0 |
| DorsAttnB |  | 2 | 5 | 7 | 6 | 6 | 9 |
| Limbic_OFC |  | 2 | 0 | 2 | 0 | 0 | 0 |
| Limbic_Pole |  | 5 | 5 | 2 | 2 | 7 | 6 |
| SalVentAttnA |  | 7 | 12 | 3 | 8 | 10 | 7 |
| SalVentAttnB |  | 1 | 7 | 1 | 0 | 2 | 6 |
| SomMotA |  | 6 | 7 | 0 | 4 | 6 | 11 |
| SomMotB |  | 2 | 2 | 0 | 0 | 0 | 3 |
| TempPar |  | 0 | 0 | 0 | 0 | 0 | 0 |
| VisCent |  | 0 | 0 | 0 | 0 | 0 | 0 |
| VisPeri |  | 0 | 5 | 3 | 2 | 3 | 1 |
| None |  | 0 | 2 | 6 | 0 | 3 | 11 |

Table S4. Counts of the cerebral cortical samples falling within each functional network of the cerebral cortical 7-network population atlas [6].

| Donors ID  Network names | | 9861 | 10021 | 12876 | 14380 | 15496 | 15697 |
| --- | --- | --- | --- | --- | --- | --- | --- |
| Control |  | 30 | 25 | 11 | 11 | 8 | 11 |
| Default |  | 74 | 58 | 30 | 41 | 42 | 33 |
| Dorsal Attention |  | 36 | 19 | 11 | 10 | 15 | 11 |
| Limbic |  | 40 | 33 | 12 | 27 | 14 | 29 |
| Ventral Attention |  | 31 | 24 | 11 | 10 | 14 | 12 |
| SomatoMotor |  | 70 | 43 | 16 | 20 | 19 | 17 |
| Visual |  | 38 | 41 | 6 | 35 | 31 | 30 |
| None |  | 147 | 118 | 78 | 101 | 71 | 97 |

Table S5. Counts of the cerebral cortical samples falling within each functional network of the cerebral cortical 17-network population atlas [6].

| Donors ID  Network names | | 9861 | 10021 | 12876 | 14380 | 15496 | 15697 |
| --- | --- | --- | --- | --- | --- | --- | --- |
| ContA |  | 11 | 15 | 8 | 2 | 6 | 4 |
| ContB |  | 22 | 2 | 6 | 0 | 5 | 5 |
| ContC |  | 2 | 1 | 4 | 6 | 4 | 3 |
| DefaultA |  | 21 | 25 | 7 | 9 | 6 | 9 |
| DefaultB |  | 26 | 19 | 14 | 22 | 21 | 17 |
| DefaultC |  | 1 | 3 | 3 | 2 | 0 | 0 |
| DorsAttnA |  | 20 | 14 | 1 | 5 | 13 | 4 |
| DorsAttnB |  | 13 | 5 | 3 | 2 | 6 | 3 |
| Limbic_OFC |  | 16 | 19 | 9 | 9 | 7 | 5 |
| Limbic_Pole |  | 24 | 15 | 3 | 20 | 9 | 14 |
| SalVentAttnA |  | 14 | 10 | 7 | 10 | 7 | 7 |
| SalVentAttnB |  | 10 | 11 | 3 | 5 | 6 | 8 |
| SomMotA |  | 34 | 24 | 9 | 10 | 11 | 14 |
| SomMotB |  | 20 | 12 | 5 | 6 | 6 | 1 |
| TempPar |  | 13 | 8 | 3 | 2 | 3 | 3 |
| VisCent |  | 12 | 13 | 0 | 16 | 11 | 9 |
| VisPeri |  | 13 | 11 | 2 | 8 | 12 | 11 |
| None |  | 194 | 154 | 88 | 121 | 81 | 113 |

Table S6. List of 59 HCP behavior measures [7].

| Category | Intuitive Name | Formal Name | Psychological Test |
| --- | --- | --- | --- |
| Alertness | MMSE | MMSE_Score | Mini Mental Status Exam |
|  | PSQI | PSQI_Score | Pittsburgh Sleep Questionnaire |
|  | TooCold | PSQI_TooCold | Pittsburgh Sleep Questionnaire |
|  | TooHot | PSQI_TooHot | Pittsburgh Sleep Questionnaire |
|  | BadDream | PSQI_BadDream | Pittsburgh Sleep Questionnaire |
|  | Pain | PSQI_Pain | Pittsburgh Sleep Questionnaire |
| Cognition | EpisodicMemory | PicSeq_AgeAdj | NIH Toolbox Picture Sequence Memory Test |
|  | CognitiveFlexibility | CardSort_AgeAdj | NIH Toolbox Dimensional Change Card Sort |
|  | Inhibition | Flanker_AgeAdj | NIH Toolbox Flanker Inhibitory Control and Attention |
|  | FluidIntelligence | PMAT24_A_CR | Penn Progressive Matrices |
|  | SkippedItems | PMAT24_A_SI | Penn Progressive Matrices |
|  | ReactionTime | PMAT24_A_RTCR | Penn Progressive Matrices |
|  | ReadingDecoding | ReadEng_AgeAdj | NIH Toolbox Oral Recognition Test |
|  | LanguageComprehension | PicVocab_AgeAdj | NIH Toolbox Picture Vocabulary Test |
|  | ProcSpeed | ProcSpeed_AgeAdj | NIH Toolbox Pattern Comparison Processing Speed Test |
|  | Impulsivity200 | DDisc_AUC_200 | Delay Discounting |
|  | Impulsivity40K | DDisc_AUC_40K | Delay Discounting |
|  | SustainedAttention-SEN | SCPT_SEN | Short Penn Continuous Performance Test |
|  | SustainedAttention-SPEC | SCPT_SPEC | Short Penn Continuous Performance Test |
|  | WorkingMemory | ListSort_AgeAdj | NIH Toolbox List Sorting Working Memory Test |
| Emotion | EmotionRecognition-CR | ER40_CR | Penn Emotion Recognition Test |
|  | EmotionRecognition-CRT | ER40_CRT | Penn Emotion Recognition Test |
|  | EmotionRecognition-Anger | ER40ANG | Penn Emotion Recognition Test |
|  | EmotionRecognition-Fear | ER40FEAR | Penn Emotion Recognition Test |
|  | EmotionRecognition-Happy | ER40HAP | Penn Emotion Recognition Test |
|  | EmotionRecognition-Neutral | ER40NOE | Penn Emotion Recognition Test |
|  | EmotionRecognition-Sad | ER40SAD | Penn Emotion Recognition Test |
|  | Anger-Affect | AngAffect_Unadj | NIH Toolbox Anger-Affect Survey |
|  | Anger-Hostility | AngHostil_Unadj | NIH Toolbox Anger-Hostility Survey |
|  | Anger-Aggression | AngAggr_Unadj | NIH Toolbox Anger-Physical Aggression Survey |
|  | Fear-Affect | FearAffect_Unadj | NIH Toolbox Fear-Affect Survey |
|  | Fear-Somatic | FearSomat_Unadj | NIH Toolbox Fear-Somatic Arousal Survey |
|  | Sadness | Sadness_Unadj | NIH Toolbox Sadness Survey |
|  | LifeSatisf | LifeSatisf_Unadj | NIH Toolbox General Life Satisfaction Survey |
|  | MeanPurp | MeanPurp_Unadj | NIH Toolbox Meaning and Purpose Survey |
|  | PosAffect | PosAffect_Unadj | NIH Toolbox Positive Affect Survey |
|  | Friendship | Friendship_Unadj Loneliness_Unadj | NIH Toolbox Friendship Survey |
|  | Loneliness | Loneliness_Unadj | NIH Toolbox Loneliness Survey |
|  | PercHostil | PercHostil_Unadj | NIH Toolbox Perceived Hostility Survey |
|  | PercReject | PercReject_Unadj | NIH Toolbox Perceived Rejection Survey |
|  | EmotSuppp | EmotSupp_Unadj | NIH Toolbox Emotional Support Survey |
|  | InstruSupp | InstruSupp_Unadj | NIH Toolbox Instrumental Support Survey |
|  | PercStress | PercStress_Unadj | NIH Toolbox Perceived Stress Survey |
|  | SelfEfficacy | SelfEff_Unadj | NIH Toolbox Self-Efficacy Survey |
| Motor | WalkEndurance | Endurance_AgeAdj | NIH Toolbox 2-minute Walk Endurance Test |
|  | GaitSpeed | GaitSpeed_Comp | NIH Toolbox 4-Meter Walk Gait Speed Test |
|  | Dexterity | Dexterity_AgeAdj | NIH Toolbox 9-hole Pegboard Dexterity Test |
|  | GripStrength | Strength_AgeAdj | NIH Toolbox Grip Strength Test |
| Personality | Agreeableness | NEOFAC_A | Five Factor Model (NEO-FFI) |
|  | Openness | NEOFAC_O | Five Factor Model (NEO-FFI) |
|  | Conscientiousness | NEOFAC_C | Five Factor Model (NEO-FFI) |
|  | Neuroticism | NEOFAC_N | Five Factor Model (NEO-FFI) |
|  | Extraversion | NEOFAC_E | Five Factor Model (NEO-FFI) |
| Sensory | Olfactory | Odor_AgeAdj | NIH Toolbox Odor Identification |
|  | PainInterf | PainInterf_Tscore | NIH Toolbox Pain Interference Survey |
|  | Gustatory | Taste_AgeAdj | NIH Toolbox Regional Taste Intensity |
|  | ContrastSensitivity-Log | Mars_Log_Score | Mars Contrast Sensitivity |
|  | ContrastSensitivity-Errs | Mars_Errs | Mars Contrast Sensitivity |
|  | ContrastSensitivity | Mars_Final | Mars Contrast Sensitivity |

Supplementary sheet 1 to sheet 28 (separate file)

Description of Supplementary sheets is described individually below.

**1. Sheet1_Gene_cere_DiffExpr**: Output of the cerebellar differential expression analyses for each cerebellar network in all 6 AHBA donors. The data in this sheet can be used to create the gene list given in "Sheet3_Gene_cere_GeneList".

**2. Sheet2_Gene_cere_Network**: Output of the cerebellar differential expression analyses for each cerebellar network in all 6 AHBA donors. The statistical threshold for differential expression is p < .05 (FDR corrected) combined with fold change > 1.

**3. Sheet3_Gene_cere_GeneList**: 443 unique genes that were positively differentially expressed across cerebellar networks in all 6 donors.

**4. Sheet4_Gene_cere_** **Specificity**: The count of overlap between differentially expressed genes in different parcellations with the 443 genes derived based on 7-network parcellation (main strategy of the present paper).

**5. Sheet5_Gene_cere_Network_4donor**: Output of the cerebellar differential expression analyses for each network in 4 left-hemisphere AHBA donors. The statistical threshold for differential expression is p < .05 (FDR corrected) combined with fold change > 1.

**6. Sheet6_Gene_cort_DiffExpr**: Output of the cortical differential expression analyses for each cerebral cortical network in all 6 AHBA donors. The data in this sheet can be used to create the gene list given in "Sheet8_Gene_cort_GeneList".

**7. Sheet7_Gene_cort_Network**: Output of the cortical differential expression analyses for each cerebral cortical network in all 6 AHBA donors. The statistical threshold for differential expression is p < .05 (FDR corrected) combined with fold change > 1.

**8. Sheet8_Gene_cort_GenenList**: 6,987 unique genes that were positively differentially expressed across cortical networks of all 6 AHBA donors.

**9. Sheet9_Gene_cortcere_Network**: Genes that are differentially expressed within both the cerebellar and cortical aspects of a given cerebello-cortical circuit, grouped by network.

**10. Sheet10_Gene_cortcere_GeneList**: 90 unique genes that are differentially expressed within the same network of both cerebellar and cortical aspects of given cerebello-cortical circuits, namely, the differentially expressed genes that overlapped within the same network of the cerebellum and the cerebral cortex.

**11. Sheet11_Gene_cere_Corr_net17**: The intra-cerebellar transcriptional correlations across 10 networks that contained samples from both 2 bi-hemisphere donors. The 10 networks were obtained based on cerebellar 17-network parcellation. These values were used to produce Figure 2A. The genes examined correspond to the 443 genes saved in "Sheet3_Gene_cere_GeneList".

**12. Sheet12_FC_cere_Corr_net17**: The intra-cerebellar resting-state functional correlations across 10 cerebellar networks that contained samples from both 2 bi-hemisphere donors. Data were from 1,018 subjects of the HCP S1200 release. These values were used to produce Figure 2B.

**13. Sheet13_Gene_cere_Corr_net7**: The intra-cerebellar transcriptional correlations across 6 networks that contained samples from both 2 bi-hemisphere donors. The 6 networks were obtained based on the cerebellar 7-network parcellation resolution. These values were used to produce Figure 2D, orange. The genes examined correspond to the 443 genes saved in "Sheet3_Gene_cere_GeneList".

**14. Sheet14_FC_cere_Corr_net7**: The intra-cerebellar resting-state functional correlations across 6 cerebellar networks that contained samples from both 2 bi-hemisphere donors. Data were from 1,018 subjects of the HCP S1200 release. These values were used to produce Figure 2D, orange.

**15. Sheet15_Gene_cere_Corr_net10**: The intra-cerebellar transcriptional correlations across 10 MDTB regions. These values were used to produce Figure 2D, blue. The genes examined correspond to the 481 genes saved in "Sheet4_Gene_cere_Specificity".

**16. Sheet16_FC_cere_Corr_net10**: The intra-cerebellar resting-state functional correlations across 10 MDTB regions. Data were from 1,018 subjects of the HCP S1200 release. These values were used to produce Figure 2D, blue.

**17. Sheet17_FC_cere_net17_unrelated**: The intra-cerebellar resting-state functional correlations across 10 cerebellar networks that contained samples from both 2 bi-hemisphere donors. Data were from 218 unrelated subjects of the HCP S1200 release.

**18. Sheet18_FC_cere_net17_HCPA**: The intra-cerebellar resting-state functional correlations across 10 cerebellar networks that contained samples from both 2 bi-hemisphere donors. Data were from 296 subjects of the HCP-Aging Lifespan 2.0 Release.

**19. Sheet19_ControlTest**: The Gene-FC correlation p values of control test based on 10,000 randomly picked 443 thresholded and thresholdless non-network-specific genes.

**20. Sheet20_Gene_cortcere_Corr**: Spearman correlation of gene expression for each cerebellar network and each of 59 cortical parcels that contained samples from both bi-hemisphere donors, using the 90 unique genes in "Sheet10_Gene_cortcere_GeneList". These values were used to produce Figure 3.

**21. Sheet21_FC_cortcere_Corr**: Pearson correlations between each cerebellar network and each of 59 cortical parcels that contained samples from both bi-hemisphere donors, using resting-state data from 1,018 subjects of the HCP S1200 release. These values were used to produce Figure 3.

**22. Sheet22_GCI+_List_n246**: The gene list of GCI^+^ (n = 246). These values were used to produce Figure 2E, left.

**23. Sheet23_GCI-_List_n197**: The gene list of GCI^-^ (n = 197). These values were used to produce Figure 2E, right.

**24. Sheet24_PCA_significant_n8**: The 8 significant principal components (PCs) derived based on the principal component analysis of the 59 behavior measures across 211 HCP unrelated participants. And the significance was evaluated by permutation test.

**25. Sheet25_Behavior-FC-Gene**: The Behavior-FC maps for PC1, PC3 and PC4 which correlated with the co-expression pattern of GCI^-^. These values were used to produce Figure 4.

**26. Sheet26_Toppgene_GCI+_n240**: The GO, pathway and disease enrichment output from ToppGene website (https://toppgene.cchmc.org/) of the 246 GCI^+^ genes. These values were used to produce Figure 5, left.

**27. Sheet27_Toppgene_GCI-_n196**: The GO, pathway and disease enrichment output from ToppGene website (https://toppgene.cchmc.org/) of the 197 GCI^-^ genes. These values were used to produce Figure 5, right.

**28. Sheet28_TemporalSpecificity**: The integrative temporal specificity expression features output from CSEA tool (http://genetics.wustl.edu/jdlab/csea-tool-2/) of GCI^+^ and GCI^-^ (FDR-BH corrected). These values were used to produce Figure 6.
